# Divergent risky decision-making and impulsivity behaviors in Lewis rat substrains with low genetic difference

**DOI:** 10.1101/2022.08.01.501451

**Authors:** Daniel B.K. Gabriel, Anna E. Liley, Hunter Franks, Monika Tutaj, Melinda R. Dwinell, Tristan de Jong, Robert W. Williams, Megan K. Mulligan, Hao Chen, Nicholas W. Simon

## Abstract

Substance use disorder (SUD) is associated with a cluster of cognitive disturbances that engender vulnerability to ongoing drug seeking and relapse. Two of these endophenotypes—risky decision-making and impulsivity—are amplified in individuals with substance use disorder and are augmented by repeated exposure to illicit drugs. Identifying genetic factors underlying variability in these traits is critical for early identification, prevention, and treatment of SUD-vulnerable individuals. Here, we compared risky decision-making and different facets of impulsivity between two fully inbred substrains of Lewis rats—LEW/NCrl and LEW/NHsd. We performed whole genome sequencing of both substrain to identify almost all relevant variants. We observed substantial differences in risky decision-making and impulsive behaviors. Relative to LEW/HHsd, the LEW/NCrl substrain accepts higher risk options in a decision-making task and higher rates of premature responses in the *differential reinforcement of low rates of responding* (DRL) task. These phenotypic differences were more pronounced in females than males. We defined a total of ∼9,000 polymorphisms between these substrains at 40X whole genome short-read coverage. Roughly half of variants are located within a single 1.5 Mb region of chromosome 8, but none impact protein-coding regions. In contrast, other variants are widely distributed, and of these 38 are predicted to cause protein-coding variants. In conclusion, Lewis rat substrains differ significantly in risk-taking and impulsivity and only a small number of easily mapped variants are likely to be causal. Sequencing combined with a reduced complexity cross (RCC) should enable identification of one or more variants underlying multiple complex addiction-relevant traits.

## Introduction

Substance use disorder (SUD) is associated with a cluster of cognitive disturbances that engender vulnerability to ongoing drug seeking and relapse ^1–5^. Two of these endophenotypes—risk-taking and impulsivity—are predictive of substance use vulnerability and are augmented by repeated drug exposure ^6–14^. Identifying the genetic factors underlying variability in these behavioral tendencies is important for understanding the molecular mechanisms driving these behaviors and is critical for early identification and ongoing treatment of SUD-vulnerable individuals.

To enable forward genetic strategies capable of identifying genetic loci and gene variants involved in risk-taking and impulsivity, it is first necessary to establish the heritability of traits in an appropriate rodent mapping population. Traditional mapping populations involve F2 intercrosses between genetically divergent inbred strains or outbred populations, both of which are segregating millions of variants. In both cases, identification of causal genes and variants responsible for trait variance is difficult. In F2 families, a low level of recombination combined with a large number of sequence variants makes it nearly impossible to identify causal QTG within each large (20 to 40 Mb) quantitative trait locus (QTL). In outbred populations, association or linkage mapping may identify a handful of candidates and associated variants or much smaller (1 to 2 Mb). However, the number of markers and individuals required for well powered mapping in outbred populations is at least 10 times higher than that needed to map using a standard F2 cross.

To address this, we quantified trait variation and heritability in substrains of rats from the inbred Lewis strain, which were developed from Wistar stock and became commercially available in the 1950s. Critically, the Lewis strain has been used in many studies of drug addiction-relevant traits ^15–17^. Rather than compare Lewis rats with another highly genetically divergent inbred strain, we obtained cohorts of Lewis rats from different breeders: Charles River Laboratories and Envigo. As both strains were initially derived from a recent common ancestor, the low genetic diversity among strains is ideal for genetic mapping using a Reduced Complexity Cross (RCC) design ^18^. RCCs are generated by intercrossing strains with highly similar genetic backgrounds to generate an F2 mapping population that segregates orders of magnitude fewer variants compared with crosses generated from more divergent strains. Large QTL intervals remain in the RCC but are no longer a barrier to QTG identification because the total number of variants is now vastly reduced–in some instances down to a single candidate gene ^19^. In addition, the low number of background variants can increase the effect of a single gene variant on trait variation, decreasing the number of individuals needed to detect a QTL ^18^. For these reasons, the RCC is an efficient strategy to identify QTGs in intercrosses derived from strains exhibiting low genetic diversity.

In the current study, we compared risky decision-making, distinct facets of impulsivity, and genetic background between Lewis rats obtained from Charles River (LEW/NCrl) and Envigo (LEW/NHsd). All subjects were obtained from their respective vendors and then bred in house for one generation to eliminate the influence of environment and shipping. Risky decision-making was measured using the RDT^20,21^, impulsive choice using the delay discounting task (DD) ^22,23^, and impulsive action using the Differential Reinforcement of Low Rate (DRL) task ^24,25^. Finally, we performed whole-genome sequencing of Lewis rats from both vendors to identify gene variants between substrains, as well as the distribution of these variants across chromosomes. The goal of these experiments was to determine if near isogenic inbred strains differ in SUD-relevant behaviors, if risk-taking and impulsivity are correlated within strain, and if these phenotypes have strong heritability for genetic mapping studies. This investigation will enable future RCC mapping studies identifying the genetic variants underlying risk-taking and impulsivity in Lewis substrains.

## Methods

### Subjects

To ensure identical breeding protocols and developmental environment, all subjects were bred in house using breeding pairs of Lewis rats obtained from either Charles River (Raleigh, NC) or Envigo (Indianapolis, IN). Charles River breeders (4 pairs) produced 27 pups (male, n = 13; female, n = 14) and Envigo breeders (4 pairs) produced 19 pups (male, n = 10; female, n = 9). At postnatal day 21, subjects were separated from dams and either pair or triple housed with subjects from their litter. Rats were housed in a 12-hour light/dark cycle (lights off at 8 a.m.), with all behavioral testing occurring during the dark cycle. When subjects reached post natal day (PND) 115, rats were food restricted to maintain 85% of free feeding weight with ad libitum access to water. All procedures were conducted in accordance with the University of Memphis Institutional Animal Care and Use Committee. While all subjects were trained in RDT, a subset of subjects did not complete delay discounting (n = 8) and DRL (n = 6) due to laboratory shutdown during the 2020 COVID-19 pandemic.

### Behavioral Apparatus

Operant chambers (Med Associates) were housed in sound attenuating cubicles. Each was equipped with a food pellet delivery trough illuminated by a 1.12 W lamp and fitted with a photobeam to detect head entries located in the center of the right wall 2 cm above the floor. On either side of the food trough there were two retractable levers located 11 cm above the floor. A 1.12 W house light was mounted on the left side wall, with a circular nose poke port equipped with a light and photo beam to detect entry directly underneath. Floors of the testing chambers consisted of steel rods connected to a shock generator that delivered scrambled foot shocks. Locomotor activity was measured during each session with infrared activity monitors located on either side of the chamber above the floor. All chambers were interfaced with a computer running custom-written codes through MedPC software (Med Associates) to control external cues and behavioral events.

### Behavioral Experiments

All subjects were trained in the 1) risky decision-making task (RDT) used to measure risk-taking, 2) delay discounting task (DD) used to measure impulsive choice, and 3) Differential Reinforcement of Low Rates of Responding (DRL) task used to measure impulsive action. To prevent task order or carryover effects, the order sequence of these three tasks was counterbalanced across subjects.

#### Instrumental Shaping

One day prior to training, all subjects were habituated in operant chambers for 5 minutes and sucrose pellet reinforcers used in behavioral experiments were placed into home cages to reduce neophobia. The following day, subjects began magazine training wherein 38 sucrose pellets were delivered to the food trough over 64 minutes. Following pellet delivery, the food trough remained illuminated until collection. Once subjects consumed the majority of pellets, they began instrumental response training.

Prior to DD or RDT, subjects learned to press a lever to receive a pellet. Upon completing 50 lever presses, subjects trained on the opposite lever. The order of levers trained was counterbalanced across subjects. After training on each lever, in a separate session subjects learned to enter the food trough when it was illuminated to cause one of the two levers to extend, which they could then press to receive a pellet reinforcer. Failure to initiate the trial or press a lever upon extension within 10s of stimulus presentation resulted in an omission. Levers were extended in a pseudorandom order until subjects completed 35 successful trials on each lever.

The final phase of shaping for either RDT or DD, reward magnitude discrimination, began with a similar design (nosepoke into lit trough to trigger lever extension), except one lever delivered a large, three pellet reward and other delivered a single pellet. After four trials of “forced choice trials” with each lever in isolation, rats were given 10 “free choice” trials in which both the small and large reward levers were extended. A press on either lever resulted in both levers retracting and delivery of the chosen reward, followed by a 10 ± 4 s second intertrial interval (ITI). Once rats displayed consistent preference for the large reward lever, they began training in either RDT or DD. Importantly, the identity of the large reward lever remained consistent across tasks.

Prior to DRL, subjects trained in a Fixed Ratio (FR)-1 protocol in which a single nosepoke into an illuminated port elicited pellet delivery. After each successful nose poke, the food trough was illuminated upon pellet delivery and remained on until subjects collected the reinforcer. The nosepoke port remained illuminated throughout the task until subjects completed 50 responses or 30 minutes had elapsed, after which the session concluded and all lights were extinguished. After receiving 50 pellets in a 30 minute session, the nose poke responses switched from an FR-1 schedule to DRL-5, and then to DRL-10 (See Methods: Impulsive Action for detailed DRL protocols).

#### Risky Decision-Making

Risky decision-making was measured using the risky decision-making task (RDT) ^26,27^. Subjects chose between a small reinforcer (one sucrose pellet) and a large reinforcer (three pellets) accompanied by a risk of foot shock (one second). Risk of foot shock increased throughout the session (0, 25, 50, 75, and 100% risk of shock) across five consecutive 18 trial blocks. Each block began with eight forced-choice trials in which a head entry to an illuminated food trough caused the lever associated with either the small or large, risky option to extend and the trough light to extinguish. Pressing this lever resulted in immediate delivery of the corresponding reinforcer, retraction of the lever, and reillumination of the food trough for 10 s or until pellet collection. The four forced-choice trials at the beginning of each block (with levers presented in pseudo-random order) enabled subjects to acquire the new risk/reward contingency. Forced choice trials were followed by ten free choice trials in which the levers associated with each outcome were presented in tandem, allowing subjects to choose between the small, safe or large, risky options. Each trial was separated by a 10 ± 4 s ITI in which the house and trough lights were extinguished and all levers were retracted. Failure to respond within 10 s of trough illumination or lever extension resulted in the trial being marked an omission, and immediate progression to the ITI. Shock intensity started at 0.05 mA, then increased in 0.05mA increments to a terminal shock level of 0.15 mA in each subsequent session. Subjects only advanced to the next shock intensity if they completed >50% of trials in the previous session. Once reaching.15mA, subjects trained in the RDT for 20 sessions, with average performance across the final five sessions used for statistical analyses.

#### Impulsive Choice

Impulsive choice was assessed through the DD task, which measures preference for small, immediate over large, delayed rewards (modified from ^22^). Each session consisted of 5 blocks with 12 trials each. Rats were presented with a choice between two levers: one lever resulted in an immediate one pellet reward, and the other was associated with a delayed 3 pellet reward, with delays of 0, 4, 8, 16, and 32s preceding pellet delivery. Each block began with two forced choice trials where either the small or large reward lever was available to establish the new delay, followed by ten free choice trials with both levers were available ^23,28,29^.

Individual trials began with illumination of the food trough and house lights. Subsequent nose poking into the food trough during this period resulted in extinguishment of the food trough light and extension of either one lever (forced choice-trials) for two trials or both levers concurrently (free choice trails) for ten trials. Failure to complete the trial initiation nose poke or lever press in the allotted time was scored as an omission. A press on either lever caused immediate retraction of both levers and immediate or delayed pellet delivery. After delivery, subjects had ten seconds to collect the pellets, then the trial proceeded to the ITI. To prevent choice of the immediate reward from reducing the time elapsed prior to the next trial, the ITI was adapted such that each trial lasted 60s regardless of delay length or choice. Subjects trained in the delay discounting for 20 sessions, and an average of behavior in the final five sessions was used for analyses.

#### Impulsive Action

To measure impulsive action, rats were trained in the DRL-10s task, in which they are required to withhold a response for a set time interval before earning a reinforcer ^30,31^. Upon completing FR-1 training, as outlined in Methods: Instrumental Shaping, subjects began DRL-5s, in which they were required to withhold a nose poke into the lit port for 5 s to receive reinforcement. Responses before this 5 s had elapsed resulted in no reinforcer delivery and the timer being reset, effectively punishing impulsive actions with a “time-out” period. After 10 sessions of DRL-5s, subjects proceeded to ten sessions of DRL-10s, in which the withholding period was increased to 10 s. Each DRL session lasted 45 minutes, with no limit on total reinforcers rats could obtain. Responses made after the withholding period had elapsed were tabulated as correct responses. Any responses prior to completion of the withholding period were scored as incorrect responses, with responses within 3 s of the previous response defined as burst responses.

#### Whole genome sequencing of Lewis Substrains

High molecular weight DNA extraction, Chromium linked reads library construction and sequencing were conducted at the HudsonAlpha Genomic Center. Spleens of one male Lewis rat from each vendor (Envigo and Charles River) were shipped to HudsonAlpha genomic center. The Qiagen MagAttract HMW DNA kit was used for DNA isolation. Sequencing library was then constructed from 1 ng of high molecular weight (∼ 50kb) genomic DNA using the Chromium Genome Library kit and sequenced on Illumina Hi-Seq (150 bp PE) using manufacturer suggested protocols.

#### Mapping of whole genome sequencing data

We obtained 878 and 863 million reads for the LEW/NCrl and LEW/NHsd substrains, respectively. Among these, 93.3% of the LEW/NCrl and 92.8% of the LEW/NHsd reads were mapped to the mRatBN7.2 reference genome. After mapping, the mean depth of coverage (i.e. the number of reads per base position for the entire genome) was 41.3X for LEW/NCrl and 40.3X for LEW/NHsd. For both strains, 2.5% of the reference genome was not present in our sequencing data (i.e., zero coverage).

#### Data Analysis

The primary variables of interest for all tasks were substrain (LEW/NCrl vs LEW/NHsd) and sex (female vs male). Average performance across the final five sessions was used as an index of performance for all three tasks, and Bivariate Pearson’s correlations were used to assess linear relationships between tasks.

For risky decision-making, percent choice of the large, risky reward in each block was used as an index of risk taking. For delay discounting, percent choice of the large, delayed reward was used as an index of impulsive choice, with less choice of the large reward defined as increased impulsive choice. In both tasks, a 3-way mixed ANOVA was conducted to identify differences based on substrain, sex, and risk level/delay. Greenhouse-Geisser corrections were used for cases in which Mauchly’s Test of Sphericity was violated; these analyses will be identifiable through presentation of non-integer degrees of freedom. Interactions were investigated using the EMMEANS and Compare subcommands within the IBM®SPSS® Statistics 26 syntax.

To provide a unitary measure of impulsive choice or risk-taking (as opposed to a multipoint curve), we utilized geometric area under the curve (AUC). AUC was calculated as the summed area of the trapezoids created by drawing vertical lines from the x-axis to percent choice of the large reinforcer in the blocks 2-5 ^32–34^. The first block was excluded from AUC calculations to maintain focus on conditions involving either a delay or risk, as the first block of each task contained no discounting factors and an objectively superior choice (3 pellets vs 1 pellet). These AUC calculations were used for correlative comparisons between tasks.

Impulsive action in DRL was calculated as correct ratio (correct choices/total choices), with increased correct ratio indicative of lower impulsive choice. In addition, for comparison between tasks (Figure 7), this measure was transformed to 1 – (correct responses/total responses), with a higher score equating to greater impulsive action. A two-way ANOVA was conducted to explore differences between the fixed factors substrain and sex as well as potential substrain x sex interactions.

Additionally, impulsive action score was compared to AUC measurements of delay discounting and risky decision-making via Pearson’s correlations. To perform a more in depth analysis of response patterns, each response in DRL was classified according to its associated inter-response-time (IRT; seconds elapsed since previous response) and plotted into a distribution consisting of total responses for 21 one second bins ^35^. All responses over 20 s were combined into a single bin and responses with an IRT < 1s were not included in analyses. A repeated-measures ANOVA with substrain and sex as between-subjects factors was used to assess effects of IRT bin on responses.

We calculated narrow-sense heritability, or the contribution of additive genetic variance to observed phenotypic variance ^36^, as a ratio of Va/(Va+Ve) ^37^, where Va is the variance of additive genetic effects and Ve is the environmental contribution to phenotypic variance. In addition to testing heritability of individual phenotypes, we created a composite risk-taking/impulsive action measure. Incorrect response ratio for DLR and % choice of risky reward were Log^10^ transformed, then we calculated the mean of these scores for each subject.

Whole genome sequencing data were mapped to the mRatBN7.2 reference genome (ref: https://wellcomeopenresearch.org/articles/6-118) using LongRanger (ver 2.2.2). DeepVariant (ver 1.0.0) was used to call SNPs and small indels from the bam files and GLnexus was used for joint calling of variants ^38^. We filtered homozygotic variant calls with PHRED quality scores greater than 30 and are unique to each substrain. SnpEff (v4_3t_core) ^39^was used for nearest gene identification and to estimate the impact of these substrain-specific variants (e.g. missense mutations, loss or gain of stop codon are classified as high impact; deletion in 5’ UTR or nonsynonymous SNP are classified as moderate impact; while synonymous variant in the coding region or start site are classified as low impact). GWAS catalog ^40^ was used to find the association of subsets of genes with psychiatric diseases or addiction related traits in human studies. GeneCup^41^ was used to explore a subset of genes for their roles in addiction or psychiatric conditions documented in the literature.

## Results

### Risky Decision-Making

We used the RDT to measure and compare risky decision-making in male and female LEW/NCrl (n=27) and LEW/NHsd (n=19) rats. Overall, subjects initially preferred the 3 pellet option, then displayed a robust shift toward the safe 1 pellet option as risk of foot shock increased (*F* (2.929, 122.999) = 40.882, *p* < 0.001; Figure 1a). Male rats showed greater risk-taking than females (*F* (1, 42) = 68.819, *p* < 0.001; Figure 1b,c), replicating previous experiments in outbred rats ^42,43^. This sex difference was observed in both substrains (*p-values* < 0.001). We also observed a block x sex interaction (*F* (2.929, 122.999) = 15.223, *p* < 0.001), such that female subjects exhibited a sharper decrease in risky choice than males in block 2, in which punishment risk was introduced (Figure 1b).

**Figure 1:**
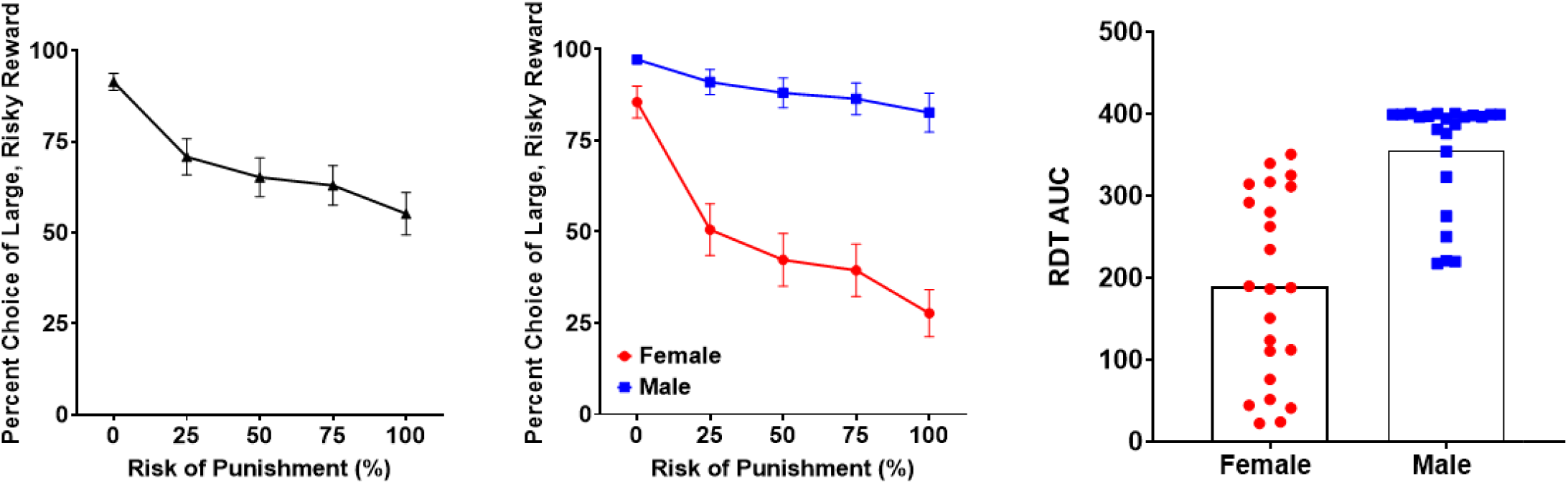
Sex differences in risky decision-making. A. Risky decision-making performance across both sexes and strains. B. Males showed greater preference for risky rewards than females. C. Individual differences in risky decision-making. Data in A-B depicted as mean±SEM.

LEW/NCrl displayed increased preference for risky options compared to LEW/NHsd (*F* _*(1, 42)*_ = 17.451, *p* < 0.001; Figure 2a), and the block x substrain interaction approached significance (*F* _*(2*.*929*, 122.99)_ = 2.437, p = 0.069). Interestingly, there was a significant substrain x sex interaction (*F* _*(1, 42)*_ = 10.1, *p* = 0.003), suggesting that the substrain differences were divergent between sexes. Pairwise comparisons revealed that female LEW/NCrl were more risk preferring than females LEW/NHsd (*p* < .001; Figure 2b), while males did not differ in risk preference between substrains (*p* = 0.477; Figure 2c). In summary, Lewis rats obtained from Charles River showed greater risk-taking than Lewis rats from Envigo, but this difference was localized to female subjects.

**Figure 2:**
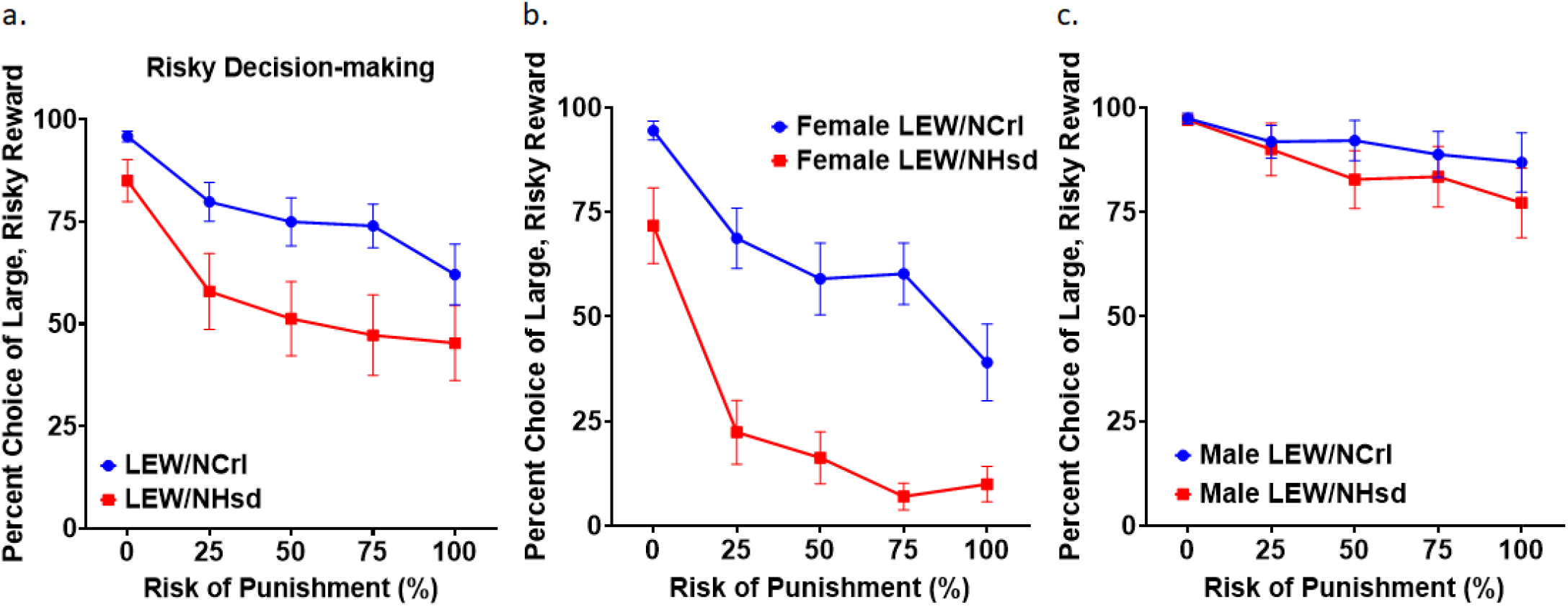
Substrain differences in risky decision-making. a. The LEW/NCrl strain showed greater risk preference than LEW/NHsd. This effect was more pronounced in females (b) than males (c).Data depicted as mean±SEM.

### Impulsive Choice

Next, we measured impulsive choice across both sexes and substrains using a delay discounting task. Subjects (n = 38) initially preferred the large reward choice, then shifting preference toward the small, immediate reward as the delay preceding the large reward increased (F _(2.086, 70.927)_ = 160.289, p < 0.001; Figure 3a). Males displayed greater choice of the large reward than females (_F (1, 34)_ = 10.102, *p* = 0.003; Figure 3b,c).

**Figure 3:**
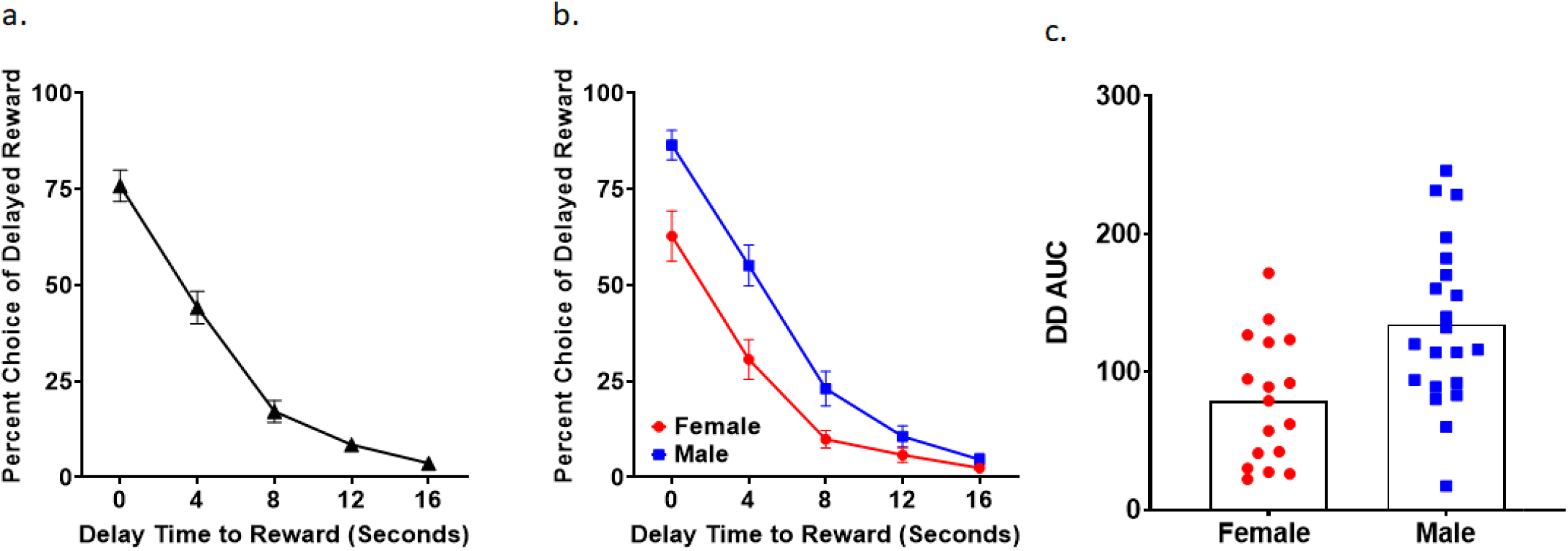
Sex differences in delay discounting. a. Delay discounting performance across all sexes/strains. b. Females showed greater delay discounting (indicative of impulsive choice) than females. c. Individual differences in delay discounting. Data in a-b depicted as mean±SEM.

There were no differences in impulsive choice between LEW/CRL and LEW/NHsd rats (effect of substrain (*F*_(1, 34)_ =1.978, *p* = .169; sex x substrain interaction: *F*_(1, 34)_ = 1.886, *p* = .179; block x substrain interaction: *F*_(2.086, 70.927)_ = .633, *p* = .541; block x sex x substrain interaction: *F* _(2.086, 70.927)_ = 1.005, *p* = .374; Figure 4). While there was no block x substrain interaction, further investigations found sex-specific substrain differences such that male LEW/NCrl discounted delayed rewards more than male LEW/NHsd (*p* = 0.032), but females did not differ between substrains (*p* = 0.983). Therefore, although on average LEW/CRL and LEW/NHsd subjects showed comparable delay discounting, there was a significant strain difference evident in males.

**Figure 4:**
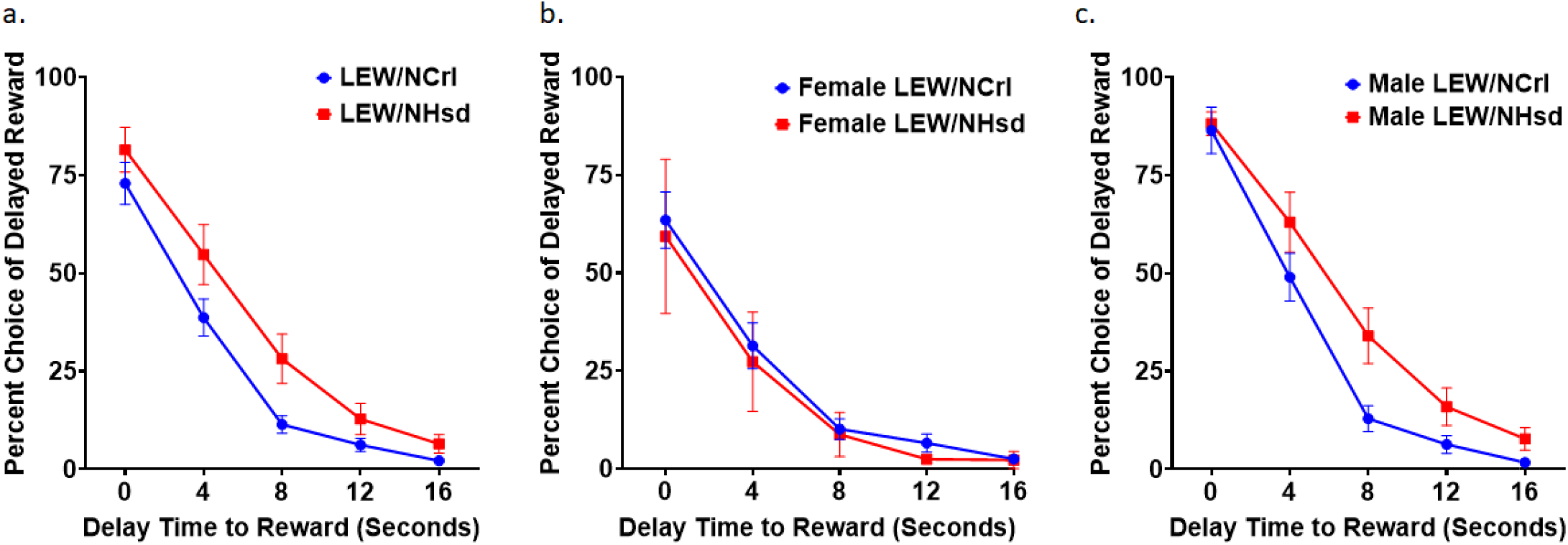
Substrain differences in delay discounting. a. No significant differences in delay discounting were observed between LEW/NCrl and LEW/NHsd strains. While there was not strain difference between females, male LEW/NCrl rats showed greater discounting than LEW/NHsd rats. All data are mean±SEM.

### Impulsive Action

Impulsive action was assessed in all subjects (n = 40; LEW/NHsd female = 12, LEW/NHsd male = 11, LEW/NCrl female = 9, LEW/NCrl male = 8) using the DRL-10s. On average, subjects were able to correctly withhold reward-seeking responses for the required 10 second period on approximately half of all responses (*M* = 50.69%, *SEM* = 1.693%). Male rats displayed a greater impulsivity score than females (*F* _*(1, 36)*_ = 6.626, *p* = 0.014; Figure 5a). LEW/NCrl had a greater impulsivity score than LEW/NHsd (*F* _*(1, 36)*_ = 4.730, *p* = 0.036), indicative of increased impulsive action (Figure 5b). There was no substrain x sex interaction (F _*(1, 36)*_ = 2.493, p = 0.123), indicating that males were more impulsive than females across both substrains (Figure 5c).

**Figure 5:**
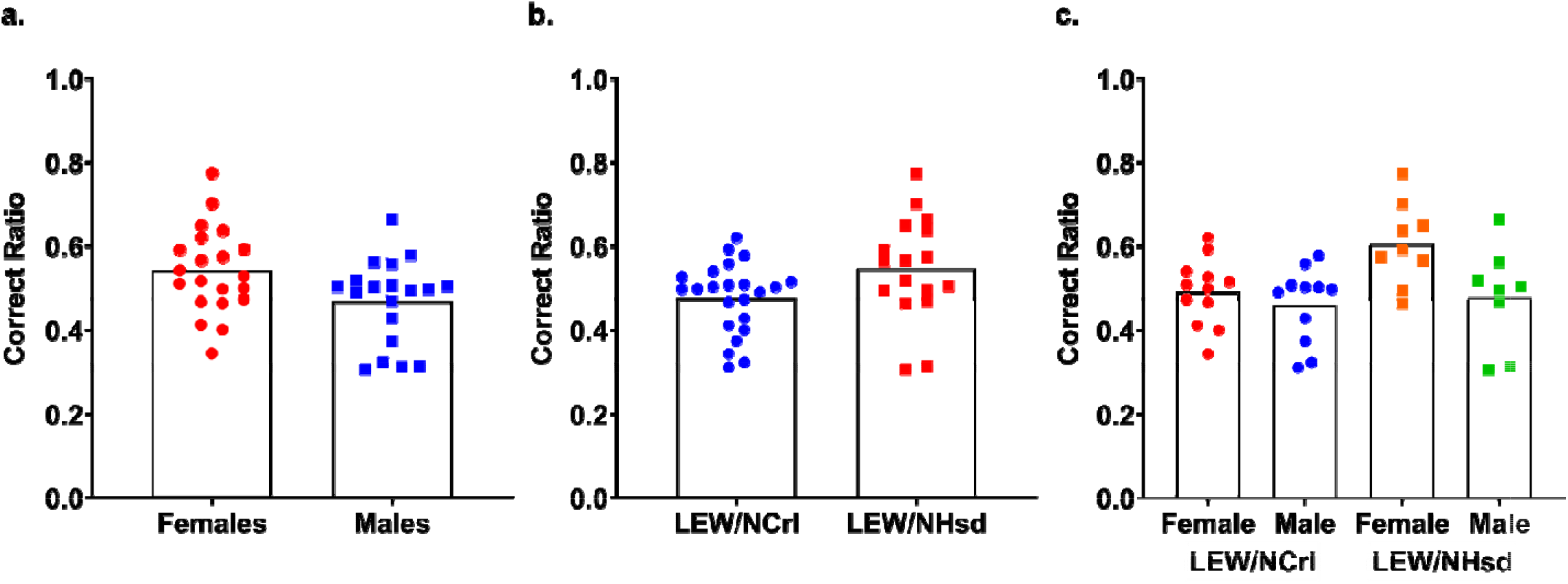
Sex and substrain differences in DRL performance. a. Females had a higher Correct Ratio in DRl-10 than males, indicating that males are more impulsive than females. b. LEW/NHsd rats had a higher Correct Ratio in DRL, indicating that LEW/NCrl are more impulsive than LEW/NHsd.

To further analyze DRL performance, total responses were divided into inter-response time (IRT) bins. As expected, this revealed a main effect of time (*F* _*(2*.*716, 97*.*759)*_ = 184.094, *p* < 0.001), with the greatest proportion of responses clustered around the 9-10 and 10-11 second bins (Figure 6a). As with impulsive action score, IRT curves differed between substrains (*F* (1, 36) = 4.419, *p* = 0.043) and sex (*F* _*(1, 36)*_ = 26.326, *p* < 0.001). When collapsed across sexes, LEW/NCrl subjects displayed an IRT curve shifted leftward from LEW/NHsd (Figure 6b) (*F* _*(2*.*716, 97*.*759)*_ = 2.810, *p* = 0.049) suggesting that these subjects were more likely to respond prior to the 10 second point than LEW/NHsd (*F* _*(1, 36)*_ = 4.419, *p* = 0.043). Therefore, the LEW/NCrl reduction in accuracy on DRL compared to LEW/NHsd was due to increased premature responses, indicative of elevated impulsive action. This effect was still observed when restricting analysis to only females (*p* = 0.008; Figure 6c), but IRT curves did not differ between substrains in males (*p* = 0.820; Figure 6d), suggesting that strain differences in impulsivity are most evident in females.

**Figure 6:**
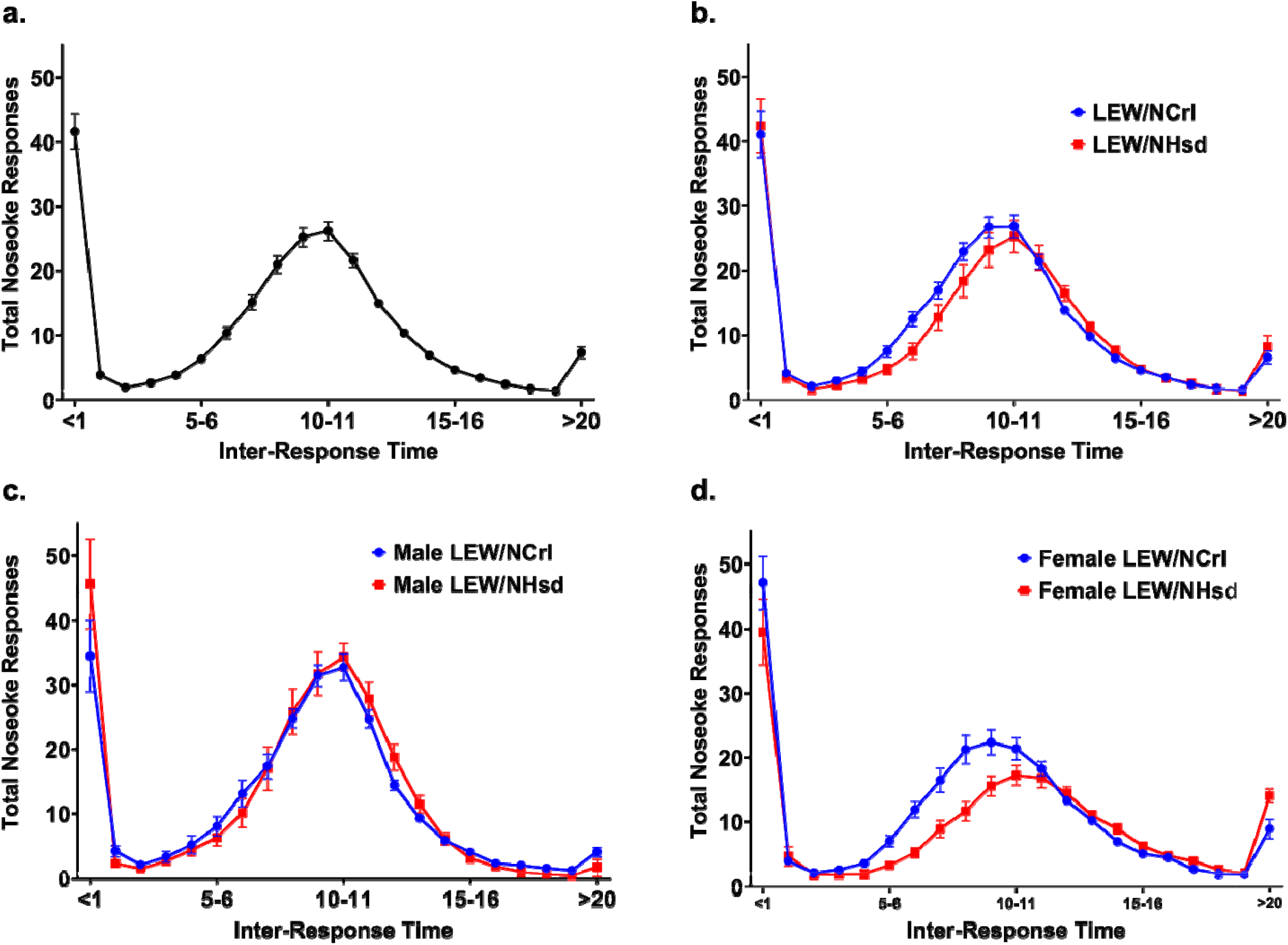
Substrain differences in inter-response time (IRT) during DRL-10. a. IRT curve displaying mean response distribution of all rats. b. IRT curve comparison between strains, with LEW/NCrl showing increased early response frequency compared to LEW/NHsd, indicative of increased impulsive action. This strain difference was more pronounced in females (c.) than males (d.). Data depicted as mean ±SEM.

### Cross Task Relationships

We compared indices of risky decision-making, impulsive choice, and impulsive action within all subjects using Pearson’s correlations. A positive correlation existed between impulsive action (1/correct ratio in DRL10) and risk-taking (AUC; *r* = -0.533, *p* < 0.001; Figure 7a). Impulsive choice, measured as delay discounting AUC, was uncorrelated with both risky decision-making (*r* = 0.177, *p* = 0.287; Figure 7b) and impulsive action (*r* = 0.058, *p* = 0.754; Figure 7c).

**Figure 7:**
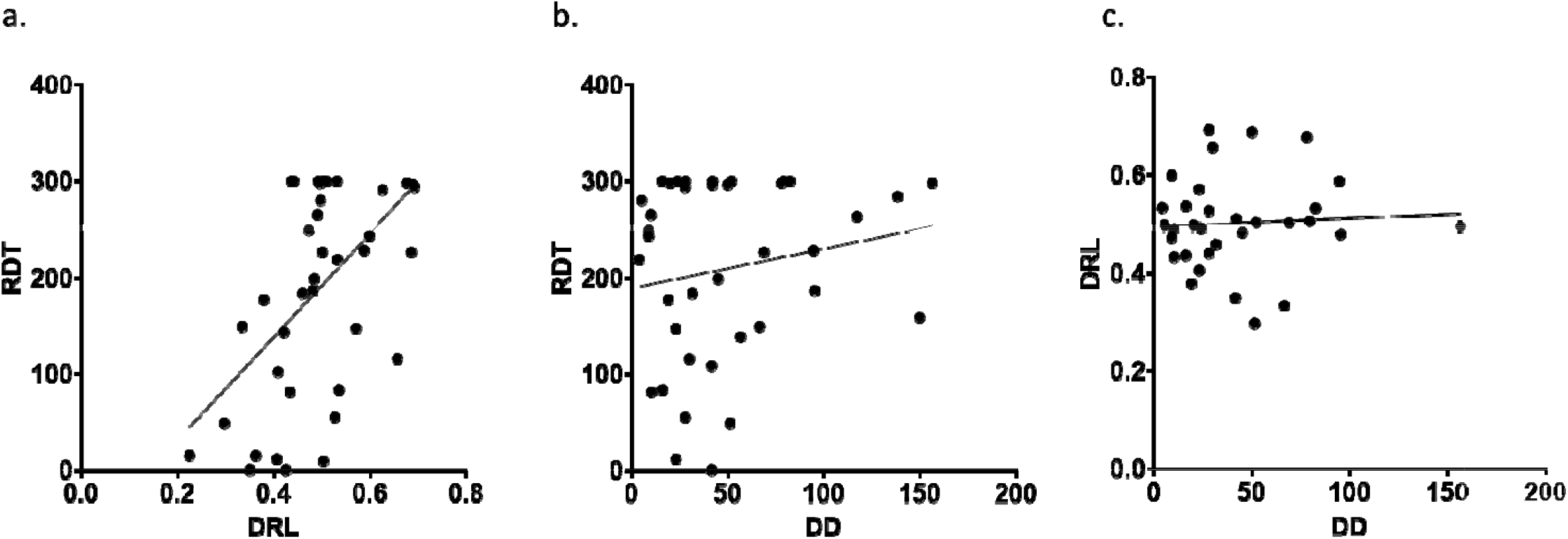
Relationships between risk-taking, impulsive choice, and impulsive action. a. There was a positive correlation between risky choice AUC and impulsivity, such that high risk preference in RDT predicted premature responding in DRL-10. There were no relationships between risk-taking and impulsive choice (b.) or impulsive action and choice (c.).

### RDT and Impulsivity Trait Heritability

Comparing LEW/NCrl with Lew/NHsd revealed high narrow-sense heritability values for each task. Heritability for risky decision-making, calculated based on AUC of risky choice, was 0.163. Heritability of impulsive action, calculated using the ratio of premature responses to total responses, was 0.148. Impulsive choice based on AUC of delayed reward choice was 0.235. Interestingly, when risky decision-making and impulsive action were merged into a composite score, the heritability was .282, suggesting that a common genetic source of variation underlies both of these phenotypes.

### Genome Sequencing

Joint variant analysis found 6,659 SNPs (2,580 in LEW/NCrl and 4,097 in LEW/NHsd) and 2,311 indels (1,017 in LEW/NCrl and 1,294 in LEW/NHsd) that differed between the two substrains. The mean genotype quality (PHRED) was 46.5 and the median was 49 for these variants. The distribution of these variants on each chromosome is shown in Figure 8 (A and B). When annotated using SnpEff, 5,268 of the total variants were located in 2,107 intergenic regions. In addition, 3,522 variants were located in the intronic regions of 1,528 genes. The classification of the remaining variants is shown in Table 1.

**Figure 8.**
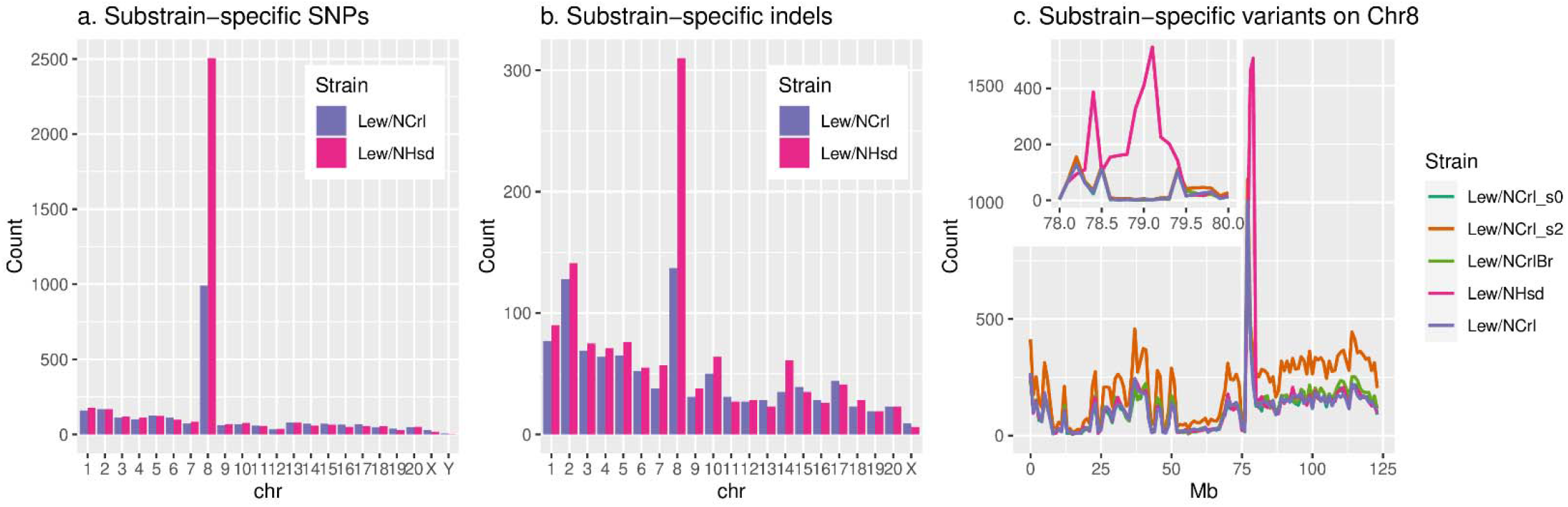
The number of SNPs and indels unique to each Lewis substrain.

**Table 1.**
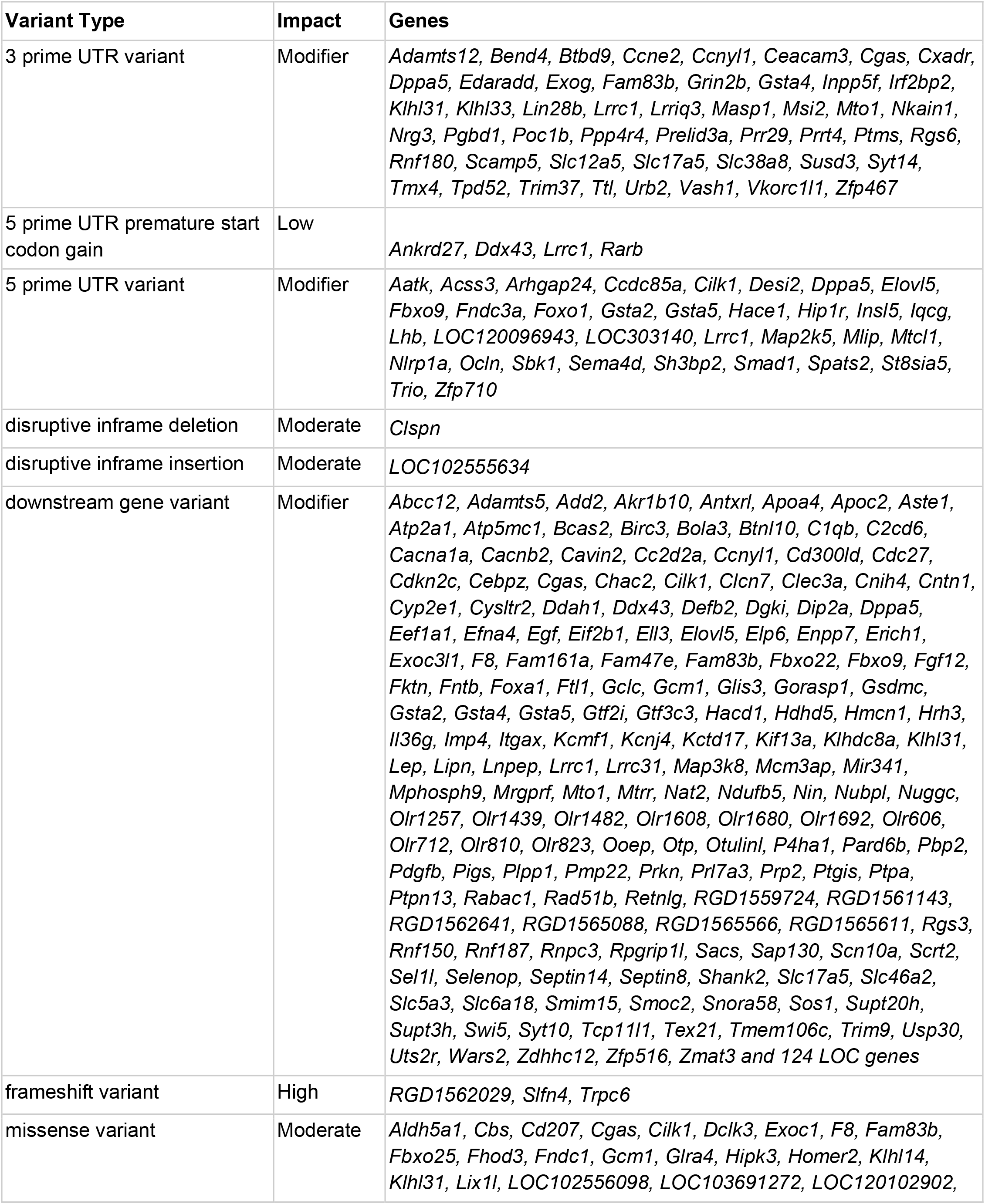
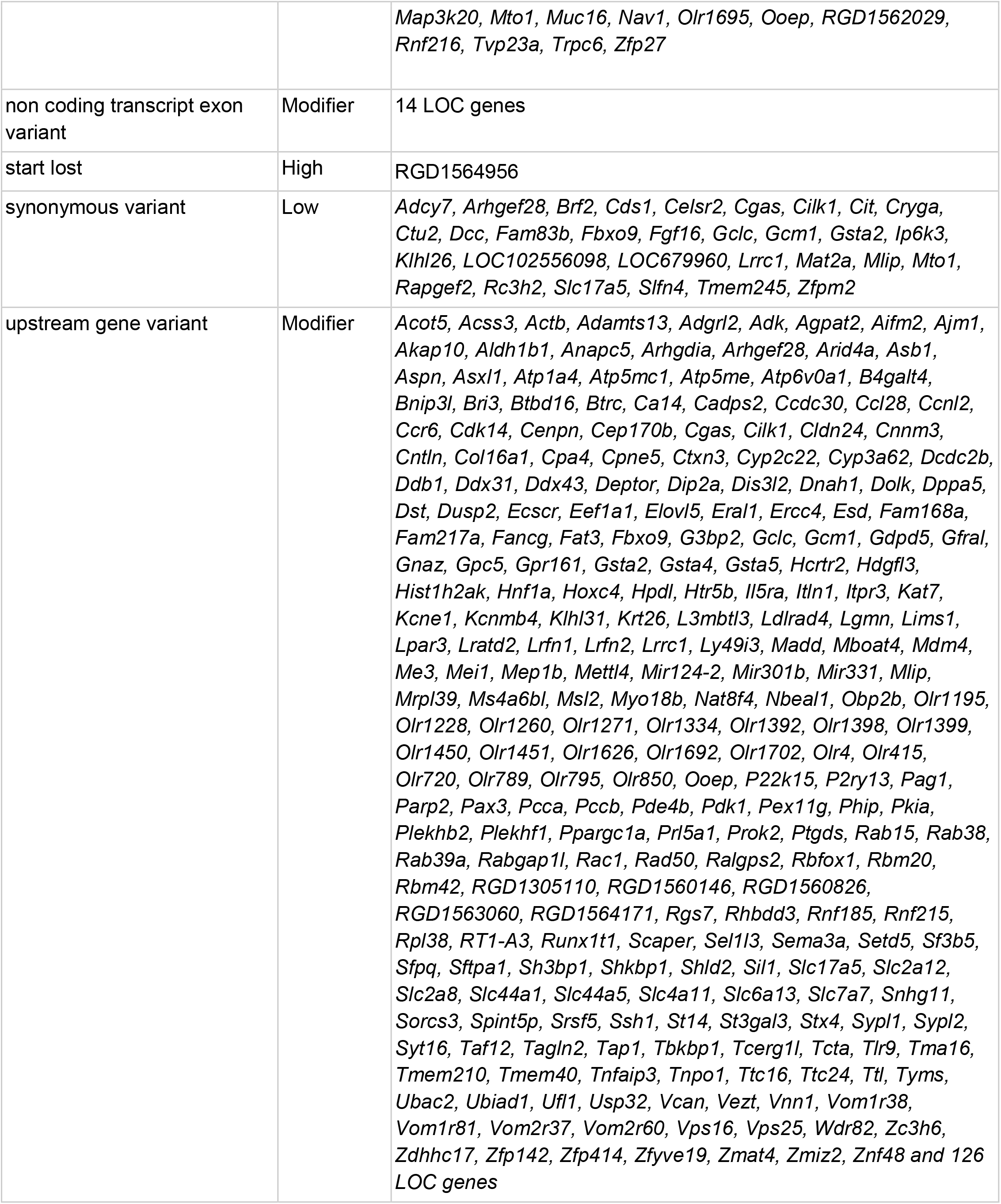
Genes with variants segregating between Lewis substrains.

Unexpectedly, chromosome (Chr) 8 contained 3,943 variants segregating between the two Lewis substrains, which accounted for 44.0% of all variants segregating them. Excluding this region, the two substrains differ by only 5,027 variants. To confirm this finding, we obtained another whole genome sequencing dataset from the Lew/NCrl substrain using Illumina WGS (Lew/NCrl_s2), and retrieved two other Lew/NCrl samples from NIH sequence read archive ^44^ (Lew/NCrl_s0 and Lew/CrlBr). As shown in Figure 8c, all four Lew/NCrl samples were consistent in the number of segregating variants on Chr 8 when compared to the Lew/NHsd sample. Further, these variants were concentrated between 78.0-79.5 Mb (Figure 8c and inset). To ensure that the substrain differences on Chr 8 were not technical in nature, we examined the genotype quality (Figure S1a), read depth (Figure S1b), and the number of missing calls (Figure S1 c) on Chr 8. These results indicated that the substrain-specific variants were unlikely to be caused by technical reasons. We also examined structural variants (SV) reported from the linked-reads dataset. Although the final call set did not contain any high-quality SV in this region, the raw barcode overlap from linked-read data, which were used for SV detection, did suggest complex SVs existed on Chr 8 in the 79.0-79.5 Mb region in both Lewis substrains (Figure S2).

### Phylogenetic Analysis

We also investigated the phylogenetic relationship between the Lewis samples and other inbred rat data found in the NIH Short Read Archive (list provided in Table S3). The phylogenetic analyses conducted on the entire Chr 8 (Figure S3a) showed that the two Lewis substrains were closely related, ruling out the possibility that our Lew/NHsd was a different strain of rat due to mistakes in sample collection. The phylogenetic analysis conducted using the genomic segment between 78.0-79.5 Mb (Figure S3b) showed that the Lew/NCrl samples were highly related to each other. Because of the lack of genetic variants in the Lew/NCrl in this region (Figure 8c, insect), they are more closely related to the reference strain (i.e., BN). In comparison, the Lew/NHsd was more closely related to the branch that contained the BN and Lew/NCrl samples than other inbred rats. Lastly, we examined the number of variants within this region on Chr 8 that were shared between the Lewis substrains and other inbred rat strains. We retained any pairs of strains that shared more than 30 variants in this region (Figure S4). We found Lew/NHsd only shared 35 variants with SBN/Ygl and SBH/Ygl strains, while Lew/NCrl shared no variants with other strains. In summary, these analyses suggested that variants in the 78-79.5 Mb regions were likely unique to the Lewis rats and were not due to technical or experimental errors. A potential explanation for the enrichment of substrain-specific variants on Chr 8 was the presence of repetitive structural variations, such as those indicated in Figure S2.

### High Impact Variants

For the variants with predicted high to moderate impact (e.g., missense or frameshift), we obtained their Gene Ontology annotation. These genes were annotated with 245 ontology categories (Table S1). We further searched genes with predicted high, moderate, or modifier impact variants in the human GWAS catalog^45^. We found 22 genes containing variants that were mapped to psychiatric or addiction related traits (Table S2).

## Discussion

These experiments revealed substantial, sex-mediated differences in addiction-relevant behavior between LEW/NHsd and LEW/NCrl substrains of rats despite high genomic similarity. Risky decision-making and impulsive action were increased in the LEW/NCrl strain, and these differences were most pronounced in female subjects. Whole genome sequencing revealed that these two substrains were only distinguished by about 9,000 genetic variants. These variants are predicted to have high to moderate impact on 38 genes/loci. Collectively, these data indicate that the LEW/NCrl strain may function as a rat model of elevated risk-taking and impulsivity. Moreover, the discovery of rat strains with divergent phenotypes despite high genetic similarity will enable future identification of the genetic basis of multiple complex addiction-relevant traits.

LEW/NHsd and LEW/NCrl strains were both derived from a common ancestor, but housed at different vendors for the past several decades (Envigo and Charles River, respectively). Therefore, strain differences may be a result of differences in handling procedures, housing, and transport to the research site. To control for experiential factors, all subjects were bred in house for one generation.

This supports the conclusion that strain differences in risk-taking and impulsivity are heritable and primarily caused by genetic variability. Importantly, these data highlight the importance of maintaining a consistent vendor within experiments or including vendor as a variable. Vendor differences in a variety of behaviors had previously been observed in outbred and Wistar rats^46–54^, and some differences were even evident between different colonies within the same vendor^55^. The current study extends this to inbred Lewis rats, and also demonstrates that phenotypic differences persist even when subjects are bred in house.

### Risky Decision-making and Impulsivity between strains

Male rats showed increased propensity for risk-taking compared to females, demonstrating that the sex differences previously observed^42,43^ in outbred rats (Long Evans) also translate to inbred strains. LEW/NCrl subjects demonstrated increased risky decision-making compared to LEW/NHsd; however, this difference was only statistically significant in females. It is possible that this sex-specificity was caused by a ceiling effect in males, as both strains approached near complete preference for the risky reward (Figure 2c). This may be a result of the relatively low shock intensity used for this experiment (0.15 mA), which is lower than the amplitude used in other experiments ^7,56–58^. As this was the first experiment (to our knowledge) to use the RDT in inbred strains, this measure was selected based on a pilot experiment which observed that higher shock amplitude evoked excessive trial omissions in female subjects. Future replications with a greater shock intensity for males may produce comparable strain differences across both sexes and increase calculated heritability. It is also possible that this strain difference is sex-dependent, suggesting that some genetic variants that mediate risk-taking may be located on sex chromosomes.

This experiment also replicated the comorbidity between risky decision-making and impulsive action observed in outbred rats^30^. This is critical because both phenotypes are commonly observed in addiction and other disorders, and both predict self-administration in rat models^6,7,59^. Risky decision-making is also predictive of cue salience and behavioral flexibility^60,61^. Therefore, future genetic mapping experiments using these rat strains may be able to detect and manipulate gene variants that drive coexpression of multiple phenotypes.

We also observed a strain difference in impulsive action. While both strains accurately achieved peak response levels at 10-11 seconds into each trial (indicative of task acquisition), LEW/NCrl rats performed a lower ratio of correct to incorrect trials than LEW/NHsd rats, and performed more premature responses prior to completion of the ten second waiting period. Males were more impulsive than females, reflected as reduced ratio of correct vs incorrect trials. As in risky decision-making, we found that the strain difference in the task was observed primarily in female subjects. We did not observe a significant difference between strains in impulsive choice (delay discounting), but did observe >0.2 narrow-sense heritability of this trait, suggesting the potential for future genetic investigation with these substrains. Surprisingly, impulsive choice had greater heritability than both risk-taking and impulsive action, despite the lack of a significant group difference. This is likely because differences in risk and impulsive action were restricted to female subjects. Finally, despite the relatively low heritability for risk-taking and impulsive action, we observed that a composite score representing both phenotypes had greater heritability than any individual behavior. This represents compelling evidence of a common source of genetic variability underlying risk-taking and impulsivity, both of which are endophenotypes of SUD, and demonstrates the feasibility of future genetic mapping experiments.

### Genetic differences between Lewis substrains

Using linked-read WGS data, we identified nearly 9,000 genetic variants that differentiated the two Lewis substrains. Surprisingly, about 44% of these strain-specific variants were located in a 1.5 Mb region (78.0 - 79.5 Mb) on Chr 8, with the remaining variants distributed across all other chromosomes. To validate this finding, we analyzed an additional previously unpublished Lew/NCrl WGS data set, and downloaded two other Lew/NCrl WGS data sets from NCBI. These three additional data sets were all in agreement with our Lew/NCrl sample (Figure 8, S1, S3, S4). To ensure that our Lew/NHsd sample was indeed from a Lewis rat, we conducted a phylogenetic analysis of Chr 8 using WGS data of 40 inbred rats. The results showed that all four Lew/NCrl samples were closely related to each other, while our Lew/NHsd sample was most similar to this collection of Lew/NCrl samples (Figure S3). Although having WGS data from a second Lew/NHsd rat is needed to confirm that our finding is applicable to all other Lew/NHsd individuals (i.e., not limited to the particular individual we selected for sequencing), the presence of similar structural features (Figure S2) between Lew/NCrl and Lew/NHsd suggested that the 1.5 Mb region on Chr 8 was initially a unique segregated feature between Lewis and other strains (e.g. Figure S4) that continued to diverge at two different breeding locations (Charles river vs Harlan/Envigo).

The genotype quality on the 1.5 Mb region with high density of variants were not different from those in adjacent regions, suggesting that it was unlikely that the quality of sequence data contributed to the substrain differences. Read depth, however, was higher in this region, from the two linked-read samples, but not in other short-read samples. Linked-read data uses molecular barcodes to restrain sequence data from the same high molecular weight DNA to be mapped next to each other, which would provide a more accurate representation of the underlying genomes. The high read depth was also validated in the barcode overlap view in this region. These data suggested the existence of relatively large (e.g. 500 Kb to 1 Mb) SVs in this region of Chr 8 in the Lewis rats. Although the SV detection pipeline did not report any high quality SV from this region, several break-ends were suggested within this region (data not shown). Future efforts using single molecule long-read sequencing techniques on additional Lew/NCrl and Lew/NHsd rats will more definitively resolve the structural differences in this Chr8 region between the Lewis substrains.

Given the collection of evidence, we believe the substrain variant hotspot on Chr 8 is likely the result of uncharacterized SVs and not due to technical artifacts or sample issues. Despite the large number of variants localized to the Chr 8 hotspot, the overall impact on risk-taking and impulsive behavior is likely to be low, This is due to the relatively low number (20) of genes/loci predicted to be impacted by variants within the hotspot, most of which have no known involvement with neuropsychiatric traits. Therefore, the phenotypic differences in risk taking and impulsivity observed between Lewis substrains is most likely caused by one or more of the ∼5,000 variants located elsewhere in the genome beyond the Chr 8 variant hotspot.

We annotated the potential impact of strain-specific genetic variants on the expression and function of genes. Using the literature mining application GeneCup^41^, we identified several of these impacted genes with known involvement in impulsivity and risk-taking behaviors and/or substance abuse. For example, *Rgs6*, which encodes the Regulator of G Protein Signaling 6 gene, contains a 3’UTR variant in the LEW/NHsd. *RGS6* variation has been associated with many psychiatric disorders associated with risk-taking and impulsivity, including obsessive-compulsive disorder, attention deficit hyperactivity disorder, schizophrenia, and bipolar disorder^62–67^. Many other variants were involved in neuronal function. For example, *Homer2*, which has a missense variant in LEW/NHsd, and *Rnf216*, which has a missense variant in LEW/NCrl, were both involved in glutamatergic synapses ^68,69^. Most relevant to this study, adolescent SHR/NCrl and Wistar rats showed decreased preference for the large but delayed reward compared with WKY/NCrl rats. This strain difference was correlated with level of *Homer2* in the prefrontal cortex of these three strains. Further, treatment with methylphenidate increased choice of the large, delayed reward, which was also accompanied by changes in mRNA levels of *Homer2*^70^. Finally, variation at the *GRIN2B* locus, which encodes the ionotropic glutamate receptor subunit 2B, has been associated with risk-tolerance and risky behavior in a GWAS with over 1 million individuals^71^. *GRIN2B* variation has also been implicated in bipolar disorder, which has elevated impulsivity and risk taking^72^ in the Chinese Han^73^ and Ashkenazi Jewish^74^ populations. Other potential genes include *MAP2K5*, which was associated with bipolar disorder^75^, and many genes, such as *LIN28B, MSI2, HIP1R* implicated in schizophrenia (Table S2).

### Summary

Our study revealed significant behavioral and genomic differences between two Lewis substrains bred in the same environment. In addition, merging risk-taking and impulsive action as a composite score yield greater heritability than either independent measure. Given the small number of genetic variants that segregate these two substrains, it is very likely that a reduced complexity cross^38^ could be used as a genetic mapping strategy to identify the causal genetic variants underlying the substrain differences in co-expressed risk taking and impulsivity.

## Notes

### Competing Interest Statement

The authors have declared no competing interest.

