## Supplemental Table 1 for "Divergent risky decision-making and impulsivity behaviors in Lewis rat substrains with low genetic difference"

| **GO** | **Gene** | **Impact** | **Relation** | **Ontology** | **Domain** |
| --- | --- | --- | --- | --- | --- |
| GO:0000076 | Clspn | disruptive_inframe_deletion | acts_upstream_of_or_within | DNA replication checkpoint signaling | biological_process |
| GO:0000077 | Map3k20 | missense_variant | involved_in | DNA damage checkpoint signaling | biological_process |
| GO:0000077 | Clspn | disruptive_inframe_deletion | involved_in | DNA damage checkpoint signaling | biological_process |
| GO:0000145 | Exoc1 | missense_variant | part_of | exocyst | cellular_component |
| GO:0001764 | Nav1 | missense_variant | involved_in | neuron migration | biological_process |
| GO:0001764 | Nav1 | missense_variant | acts_upstream_of_or_within | neuron migration | biological_process |
| GO:0001933 | Klhl31 | missense_variant | involved_in | negative regulation of protein phosphorylation | biological_process |
| GO:0001958 | Cbs | missense_variant | acts_upstream_of_or_within | endochondral ossification | biological_process |
| GO:0001974 | Cbs | missense_variant | acts_upstream_of_or_within | blood vessel remodeling | biological_process |
| GO:0002098 | Mto1 | missense_variant | involved_in | tRNA wobble uridine modification | biological_process |
| GO:0002098 | Mto1 | missense_variant | acts_upstream_of_or_within | tRNA wobble uridine modification | biological_process |
| GO:0003674 | Muc16 | missense_variant | enables | molecular_function | molecular_function |
| GO:0003674 | Med15 | splice_donor_variant&intron_variant | enables | molecular_function | molecular_function |
| GO:0003674 | Lix1l | missense_variant | enables | molecular_function | molecular_function |
| GO:0003677 | Gcm1 | missense_variant | enables | DNA binding | molecular_function |
| GO:0003677 | Cgas | missense_variant | enables | DNA binding | molecular_function |
| GO:0003677 | Hipk3 | missense_variant | enables | DNA binding | molecular_function |
| GO:0003682 | Cgas | missense_variant | enables | chromatin binding | molecular_function |
| GO:0003690 | Cgas | missense_variant | enables | double-stranded DNA binding | molecular_function |
| GO:0003700 | Gcm1 | missense_variant | enables | DNA-binding transcription factor activity | molecular_function |
| GO:0003712 | Med15 | splice_donor_variant&intron_variant | enables | transcription coregulator activity | molecular_function |
| GO:0003723 | Ooep | missense_variant | enables | RNA binding | molecular_function |
| GO:0003729 | Ooep | missense_variant | enables | mRNA binding | molecular_function |
| GO:0004672 | Hipk3 | missense_variant | enables | protein kinase activity | molecular_function |
| GO:0004672 | Map3k20 | missense_variant | enables | protein kinase activity | molecular_function |
| GO:0004672 | Dclk3 | missense_variant | enables | protein kinase activity | molecular_function |
| GO:0004672 | Cilk1 | missense_variant | enables | protein kinase activity | molecular_function |
| GO:0004674 | Cilk1 | missense_variant | enables | protein serine/threonine kinase activity | molecular_function |
| GO:0004674 | Hipk3 | missense_variant | enables | protein serine/threonine kinase activity | molecular_function |
| GO:0004674 | Map3k20 | missense_variant | enables | protein serine/threonine kinase activity | molecular_function |
| GO:0004674 | Hipk3 | missense_variant | enables | protein serine/threonine kinase activity | molecular_function |
| GO:0004674 | Cilk1 | missense_variant | enables | protein serine/threonine kinase activity | molecular_function |
| GO:0004709 | Map3k20 | missense_variant | enables | MAP kinase kinase kinase activity | molecular_function |
| GO:0004709 | Map3k20 | missense_variant | enables | MAP kinase kinase kinase activity | molecular_function |
| GO:0004713 | Hipk3 | missense_variant | enables | protein tyrosine kinase activity | molecular_function |
| GO:0004777 | Aldh5a1 | missense_variant | enables | succinate-semialdehyde dehydrogenase (NAD+) activity | molecular_function |
| GO:0004777 | Aldh5a1 | missense_variant | enables | succinate-semialdehyde dehydrogenase (NAD+) activity | molecular_function |
| GO:0006468 | Map3k20 | missense_variant | involved_in | protein phosphorylation | biological_process |
| GO:0006468 | Cilk1 | missense_variant | involved_in | protein phosphorylation | biological_process |
| GO:0006468 | Ooep | missense_variant | acts_upstream_of_or_within | protein phosphorylation | biological_process |
| GO:0006468 | Hipk3 | missense_variant | involved_in | protein phosphorylation | biological_process |
| GO:0006535 | Cbs | missense_variant | involved_in | cysteine biosynthetic process from serine | biological_process |
| GO:0006536 | Aldh5a1 | missense_variant | involved_in | glutamate metabolic process | biological_process |
| GO:0006536 | Aldh5a1 | missense_variant | acts_upstream_of_or_within | glutamate metabolic process | biological_process |
| GO:0006563 | Cbs | missense_variant | involved_in | L-serine metabolic process | biological_process |
| GO:0006565 | Cbs | missense_variant | involved_in | L-serine catabolic process | biological_process |
| GO:0006801 | Cbs | missense_variant | acts_upstream_of_or_within | superoxide metabolic process | biological_process |
| GO:0006828 | Trpc6 | frameshift_variant | involved_in | manganese ion transport | biological_process |
| GO:0006828 | Trpc6 | missense_variant | involved_in | manganese ion transport | biological_process |
| GO:0006887 | Exoc1 | missense_variant | involved_in | exocytosis | biological_process |
| GO:0006893 | Exoc1 | missense_variant | involved_in | Golgi to plasma membrane transport | biological_process |
| GO:0006915 | Hipk3 | missense_variant | involved_in | apoptotic process | biological_process |
| GO:0006974 | Cgas | missense_variant | involved_in | cellular response to DNA damage stimulus | biological_process |
| GO:0006974 | Cgas | missense_variant | involved_in | cellular response to DNA damage stimulus | biological_process |
| GO:0007010 | Map3k20 | missense_variant | acts_upstream_of_or_within | cytoskeleton organization | biological_process |
| GO:0007015 | Ooep | missense_variant | involved_in | actin filament organization | biological_process |
| GO:0007095 | Clspn | disruptive_inframe_deletion | involved_in | mitotic G2 DNA damage checkpoint signaling | biological_process |
| GO:0007095 | Clspn | disruptive_inframe_deletion | involved_in | mitotic G2 DNA damage checkpoint signaling | biological_process |
| GO:0007165 | Glra4 | missense_variant | involved_in | signal transduction | biological_process |
| GO:0007165 | Fam83b | missense_variant | involved_in | signal transduction | biological_process |
| GO:0007173 | Fam83b | missense_variant | involved_in | epidermal growth factor receptor signaling pathway | biological_process |
| GO:0007173 | Fam83b | missense_variant | involved_in | epidermal growth factor receptor signaling pathway | biological_process |
| GO:0007186 | Olr1695 | missense_variant | involved_in | G protein-coupled receptor signaling pathway | biological_process |
| GO:0007204 | Trpc6 | missense_variant | involved_in | positive regulation of cytosolic calcium ion concentration | biological_process |
| GO:0007204 | Trpc6 | missense_variant | acts_upstream_of_or_within | positive regulation of cytosolic calcium ion concentration | biological_process |
| GO:0007204 | Trpc6 | frameshift_variant | involved_in | positive regulation of cytosolic calcium ion concentration | biological_process |
| GO:0007204 | Trpc6 | frameshift_variant | acts_upstream_of_or_within | positive regulation of cytosolic calcium ion concentration | biological_process |
| GO:0007216 | Homer2 | missense_variant | involved_in | G protein-coupled glutamate receptor signaling pathway | biological_process |
| GO:0007218 | Glra4 | missense_variant | involved_in | neuropeptide signaling pathway | biological_process |
| GO:0007254 | Map3k20 | missense_variant | involved_in | JNK cascade | biological_process |
| GO:0007268 | Glra4 | missense_variant | involved_in | chemical synaptic transmission | biological_process |
| GO:0007338 | Trpc6 | frameshift_variant | involved_in | single fertilization | biological_process |
| GO:0007338 | Trpc6 | missense_variant | involved_in | single fertilization | biological_process |
| GO:0007399 | Nav1 | missense_variant | involved_in | nervous system development | biological_process |
| GO:0007417 | Aldh5a1 | missense_variant | involved_in | central nervous system development | biological_process |
| GO:0007566 | Exoc1 | missense_variant | acts_upstream_of_or_within | embryo implantation | biological_process |
| GO:0007566 | Ooep | missense_variant | acts_upstream_of_or_within | embryo implantation | biological_process |
| GO:0007568 | Trpc6 | frameshift_variant | involved_in | aging | biological_process |
| GO:0007568 | Trpc6 | missense_variant | involved_in | aging | biological_process |
| GO:0007596 | F8 | missense_variant | involved_in | blood coagulation | biological_process |
| GO:0007597 | F8 | missense_variant | involved_in | blood coagulation, intrinsic pathway | biological_process |
| GO:0007597 | F8 | missense_variant | involved_in | blood coagulation, intrinsic pathway | biological_process |
| GO:0007605 | Homer2 | missense_variant | involved_in | sensory perception of sound | biological_process |
| GO:0008150 | Med15 | splice_donor_variant&intron_variant | involved_in | biological_process | biological_process |
| GO:0008150 | Lix1l | missense_variant | involved_in | biological_process | biological_process |
| GO:0008150 | Muc16 | missense_variant | involved_in | biological_process | biological_process |
| GO:0008270 | Gcm1 | missense_variant | enables | zinc ion binding | molecular_function |
| GO:0008277 | Homer2 | missense_variant | acts_upstream_of_or_within | regulation of G protein-coupled receptor signaling pathway | biological_process |
| GO:0008283 | Fam83b | missense_variant | involved_in | cell population proliferation | biological_process |
| GO:0008340 | Cgas | missense_variant | acts_upstream_of_or_within | determination of adult lifespan | biological_process |
| GO:0010468 | Ooep | missense_variant | involved_in | regulation of gene expression | biological_process |
| GO:0010666 | Fndc1 | missense_variant | acts_upstream_of_or_within | positive regulation of cardiac muscle cell apoptotic process | biological_process |
| GO:0010749 | Cbs | missense_variant | acts_upstream_of_or_within | regulation of nitric oxide mediated signal transduction | biological_process |
| GO:0010753 | Cgas | missense_variant | involved_in | positive regulation of cGMP-mediated signaling | biological_process |
| GO:0010753 | Cgas | missense_variant | involved_in | positive regulation of cGMP-mediated signaling | biological_process |
| GO:0010800 | Trpc6 | missense_variant | involved_in | positive regulation of peptidyl-threonine phosphorylation | biological_process |
| GO:0010800 | Trpc6 | frameshift_variant | involved_in | positive regulation of peptidyl-threonine phosphorylation | biological_process |
| GO:0010997 | Clspn | disruptive_inframe_deletion | enables | anaphase-promoting complex binding | molecular_function |
| GO:0010997 | Clspn | disruptive_inframe_deletion | enables | anaphase-promoting complex binding | molecular_function |
| GO:0014069 | Homer2 | missense_variant | located_in | postsynaptic density | cellular_component |
| GO:0014069 | Homer2 | missense_variant | is_active_in | postsynaptic density | cellular_component |
| GO:0014069 | Homer2 | missense_variant | is_active_in | postsynaptic density | cellular_component |
| GO:0015279 | Trpc6 | missense_variant | enables | store-operated calcium channel activity | molecular_function |
| GO:0015279 | Trpc6 | frameshift_variant | enables | store-operated calcium channel activity | molecular_function |
| GO:0015279 | Trpc6 | missense_variant | enables | store-operated calcium channel activity | molecular_function |
| GO:0015279 | Trpc6 | frameshift_variant | enables | store-operated calcium channel activity | molecular_function |
| GO:0015629 | Klhl14 | missense_variant | located_in | actin cytoskeleton | cellular_component |
| GO:0015630 | Nav1 | missense_variant | is_active_in | microtubule cytoskeleton | cellular_component |
| GO:0015630 | Nav1 | missense_variant | located_in | microtubule cytoskeleton | cellular_component |
| GO:0016491 | F8 | missense_variant | enables | oxidoreductase activity | molecular_function |
| GO:0016567 | Fbxo25 | missense_variant | involved_in | protein ubiquitination | biological_process |
| GO:0016567 | Fbxo25 | missense_variant | involved_in | protein ubiquitination | biological_process |
| GO:0016567 | Fbxo25 | missense_variant | involved_in | protein ubiquitination | biological_process |
| GO:0016592 | Med15 | splice_donor_variant&intron_variant | part_of | mediator complex | cellular_component |
| GO:0016594 | Glra4 | missense_variant | contributes_to | glycine binding | molecular_function |
| GO:0016604 | Hipk3 | missense_variant | located_in | nuclear body | cellular_component |
| GO:0016605 | Hipk3 | missense_variant | is_active_in | PML body | cellular_component |
| GO:0016605 | Hipk3 | missense_variant | located_in | PML body | cellular_component |
| GO:0016607 | Fndc1 | missense_variant | located_in | nuclear speck | cellular_component |
| GO:0016934 | Glra4 | missense_variant | contributes_to | extracellularly glycine-gated chloride channel activity | molecular_function |
| GO:0016934 | Glra4 | missense_variant | enables | extracellularly glycine-gated chloride channel activity | molecular_function |
| GO:0019825 | Cbs | missense_variant | enables | oxygen binding | molecular_function |
| GO:0019827 | Med15 | splice_donor_variant&intron_variant | acts_upstream_of_or_within | stem cell population maintenance | biological_process |
| GO:0019898 | Muc16 | missense_variant | located_in | extrinsic component of membrane | cellular_component |
| GO:0019899 | Cbs | missense_variant | enables | enzyme binding | molecular_function |
| GO:0019901 | Fam83b | missense_variant | enables | protein kinase binding | molecular_function |
| GO:0019901 | Fam83b | missense_variant | enables | protein kinase binding | molecular_function |
| GO:0019904 | Homer2 | missense_variant | enables | protein domain specific binding | molecular_function |
| GO:0020037 | Cbs | missense_variant | enables | heme binding | molecular_function |
| GO:0021587 | Cbs | missense_variant | acts_upstream_of_or_within | cerebellum morphogenesis | biological_process |
| GO:0022824 | Glra4 | missense_variant | enables | transmitter-gated ion channel activity | molecular_function |
| GO:0030017 | Fhod3 | missense_variant | located_in | sarcomere | cellular_component |
| GO:0030160 | Homer2 | missense_variant | enables | synaptic receptor adaptor activity | molecular_function |
| GO:0030170 | Cbs | missense_variant | enables | pyridoxal phosphate binding | molecular_function |
| GO:0030173 | Tvp23a | missense_variant&splice_region_variant | is_active_in | integral component of Golgi membrane | cellular_component |
| GO:0030182 | Trpc6 | missense_variant | involved_in | neuron differentiation | biological_process |
| GO:0030182 | Trpc6 | frameshift_variant | involved_in | neuron differentiation | biological_process |
| GO:0030246 | Cd207 | missense_variant | enables | carbohydrate binding | molecular_function |
| GO:0030276 | Trpc6 | missense_variant | enables | clathrin binding | molecular_function |
| GO:0030276 | Trpc6 | frameshift_variant | enables | clathrin binding | molecular_function |
| GO:0030425 | Homer2 | missense_variant | is_active_in | dendrite | cellular_component |
| GO:0030425 | Homer2 | missense_variant | located_in | dendrite | cellular_component |
| GO:0030488 | Mto1 | missense_variant | involved_in | tRNA methylation | biological_process |
| GO:0030837 | Fhod3 | missense_variant | acts_upstream_of_or_within | negative regulation of actin filament polymerization | biological_process |
| GO:0030866 | Fhod3 | missense_variant | involved_in | cortical actin cytoskeleton organization | biological_process |
| GO:0031297 | Ooep | missense_variant | involved_in | replication fork processing | biological_process |
| GO:0031406 | Aldh5a1 | missense_variant | enables | carboxylic acid binding | molecular_function |
| GO:0031966 | Fndc1 | missense_variant | located_in | mitochondrial membrane | cellular_component |
| GO:0032147 | Clspn | disruptive_inframe_deletion | involved_in | activation of protein kinase activity | biological_process |
| GO:0032147 | Clspn | disruptive_inframe_deletion | involved_in | activation of protein kinase activity | biological_process |
| GO:0032414 | Trpc6 | frameshift_variant | acts_upstream_of_or_within | positive regulation of ion transmembrane transporter activity | biological_process |
| GO:0032414 | Trpc6 | missense_variant | acts_upstream_of_or_within | positive regulation of ion transmembrane transporter activity | biological_process |
| GO:0032426 | Homer2 | missense_variant | located_in | stereocilium tip | cellular_component |
| GO:0032479 | Cgas | missense_variant | acts_upstream_of_or_within | regulation of type I interferon production | biological_process |
| GO:0032481 | Cgas | missense_variant | involved_in | positive regulation of type I interferon production | biological_process |
| GO:0032481 | Cgas | missense_variant | involved_in | positive regulation of type I interferon production | biological_process |
| GO:0032648 | Rnf216 | missense_variant | involved_in | regulation of interferon-beta production | biological_process |
| GO:0032703 | Homer2 | missense_variant | involved_in | negative regulation of interleukin-2 production | biological_process |
| GO:0032715 | Muc16 | missense_variant | acts_upstream_of_or_within | negative regulation of interleukin-6 production | biological_process |
| GO:0032880 | Ooep | missense_variant | involved_in | regulation of protein localization | biological_process |
| GO:0032991 | Ooep | missense_variant | part_of | protein-containing complex | cellular_component |
| GO:0033314 | Clspn | disruptive_inframe_deletion | involved_in | mitotic DNA replication checkpoint signaling | biological_process |
| GO:0033314 | Clspn | disruptive_inframe_deletion | involved_in | mitotic DNA replication checkpoint signaling | biological_process |
| GO:0035088 | Ooep | missense_variant | acts_upstream_of_or_within | establishment or maintenance of apical/basal cell polarity | biological_process |
| GO:0035254 | Homer2 | missense_variant | enables | glutamate receptor binding | molecular_function |
| GO:0035256 | Homer2 | missense_variant | enables | G protein-coupled glutamate receptor binding | molecular_function |
| GO:0035256 | Homer2 | missense_variant | enables | G protein-coupled glutamate receptor binding | molecular_function |
| GO:0035556 | Dclk3 | missense_variant | involved_in | intracellular signal transduction | biological_process |
| GO:0035556 | Map3k20 | missense_variant | involved_in | intracellular signal transduction | biological_process |
| GO:0035556 | Cilk1 | missense_variant | involved_in | intracellular signal transduction | biological_process |
| GO:0035556 | Cilk1 | missense_variant | involved_in | intracellular signal transduction | biological_process |
| GO:0035584 | Homer2 | missense_variant | acts_upstream_of_or_within | calcium-mediated signaling using intracellular calcium source | biological_process |
| GO:0035720 | Cilk1 | missense_variant | involved_in | intraciliary anterograde transport | biological_process |
| GO:0035721 | Cilk1 | missense_variant | involved_in | intraciliary retrograde transport | biological_process |
| GO:0035861 | Cgas | missense_variant | is_active_in | site of double-strand break | cellular_component |
| GO:0035861 | Cgas | missense_variant | located_in | site of double-strand break | cellular_component |
| GO:0038001 | Cgas | missense_variant | involved_in | paracrine signaling | biological_process |
| GO:0038001 | Cgas | missense_variant | involved_in | paracrine signaling | biological_process |
| GO:0038066 | Map3k20 | missense_variant | involved_in | p38MAPK cascade | biological_process |
| GO:0042802 | Aldh5a1 | missense_variant | enables | identical protein binding | molecular_function |
| GO:0042802 | Cbs | missense_variant | enables | identical protein binding | molecular_function |
| GO:0042802 | Homer2 | missense_variant | enables | identical protein binding | molecular_function |
| GO:0042803 | Cbs | missense_variant | enables | protein homodimerization activity | molecular_function |
| GO:0042803 | Trpc6 | frameshift_variant | enables | protein homodimerization activity | molecular_function |
| GO:0042803 | Trpc6 | missense_variant | enables | protein homodimerization activity | molecular_function |
| GO:0042805 | Trpc6 | frameshift_variant | enables | actinin binding | molecular_function |
| GO:0042805 | Trpc6 | missense_variant | enables | actinin binding | molecular_function |
| GO:0042826 | Gcm1 | missense_variant | enables | histone deacetylase binding | molecular_function |
| GO:0043005 | Klhl14 | missense_variant | is_active_in | neuron projection | cellular_component |
| GO:0043005 | Glra4 | missense_variant | is_active_in | neuron projection | cellular_component |
| GO:0043005 | Klhl14 | missense_variant | located_in | neuron projection | cellular_component |
| GO:0043025 | Klhl14 | missense_variant | is_active_in | neuronal cell body | cellular_component |
| GO:0043025 | Klhl14 | missense_variant | located_in | neuronal cell body | cellular_component |
| GO:0043025 | Homer2 | missense_variant | located_in | neuronal cell body | cellular_component |
| GO:0043065 | Map3k20 | missense_variant | involved_in | positive regulation of apoptotic process | biological_process |
| GO:0043066 | Cbs | missense_variant | involved_in | negative regulation of apoptotic process | biological_process |
| GO:0043066 | Hipk3 | missense_variant | involved_in | negative regulation of apoptotic process | biological_process |
| GO:0043418 | Cbs | missense_variant | involved_in | homocysteine catabolic process | biological_process |
| GO:0043506 | Cbs | missense_variant | acts_upstream_of_or_within | regulation of JUN kinase activity | biological_process |
| GO:0043508 | Hipk3 | missense_variant | involved_in | negative regulation of JUN kinase activity | biological_process |
| GO:0043950 | Cgas | missense_variant | involved_in | positive regulation of cAMP-mediated signaling | biological_process |
| GO:0043950 | Cgas | missense_variant | involved_in | positive regulation of cAMP-mediated signaling | biological_process |
| GO:0048015 | Exoc1 | missense_variant | involved_in | phosphatidylinositol-mediated signaling | biological_process |
| GO:0048286 | Slfn4 | frameshift_variant | involved_in | lung alveolus development | biological_process |
| GO:0050421 | Cbs | missense_variant | enables | nitrite reductase (NO-forming) activity | molecular_function |
| GO:0050660 | Mto1 | missense_variant | enables | flavin adenine dinucleotide binding | molecular_function |
| GO:0050667 | Cbs | missense_variant | acts_upstream_of_or_within | homocysteine metabolic process | biological_process |
| GO:0050667 | Cbs | missense_variant | involved_in | homocysteine metabolic process | biological_process |
| GO:0050680 | Muc16 | missense_variant | acts_upstream_of_or_within | negative regulation of epithelial cell proliferation | biological_process |
| GO:0050691 | Rnf216 | missense_variant | involved_in | regulation of defense response to virus by host | biological_process |
| GO:0050714 | Exoc1 | missense_variant | involved_in | positive regulation of protein secretion | biological_process |
| GO:0051593 | Cbs | missense_variant | acts_upstream_of_or_within | response to folic acid | biological_process |
| GO:0051607 | Cgas | missense_variant | involved_in | defense response to virus | biological_process |
| GO:0051607 | Cd207 | missense_variant | involved_in | defense response to virus | biological_process |
| GO:0051639 | Fhod3 | missense_variant | acts_upstream_of_or_within | actin filament network formation | biological_process |
| GO:0051639 | Fhod3 | missense_variant | involved_in | actin filament network formation | biological_process |
| GO:0051726 | Map3k20 | missense_variant | involved_in | regulation of cell cycle | biological_process |
| GO:0051928 | Trpc6 | frameshift_variant | acts_upstream_of_or_within | positive regulation of calcium ion transport | biological_process |
| GO:0051928 | Trpc6 | missense_variant | acts_upstream_of_or_within | positive regulation of calcium ion transport | biological_process |
| GO:0055003 | Fhod3 | missense_variant | acts_upstream_of_or_within | cardiac myofibril assembly | biological_process |
| GO:0055003 | Fhod3 | missense_variant | involved_in | cardiac myofibril assembly | biological_process |
| GO:0070301 | Trpc6 | frameshift_variant | involved_in | cellular response to hydrogen peroxide | biological_process |
| GO:0070301 | Trpc6 | missense_variant | involved_in | cellular response to hydrogen peroxide | biological_process |
| GO:0070679 | Trpc6 | frameshift_variant | enables | inositol 1,4,5 trisphosphate binding | molecular_function |
| GO:0070679 | Trpc6 | frameshift_variant | enables | inositol 1,4,5 trisphosphate binding | molecular_function |
| GO:0070679 | Trpc6 | missense_variant | enables | inositol 1,4,5 trisphosphate binding | molecular_function |
| GO:0070679 | Trpc6 | missense_variant | enables | inositol 1,4,5 trisphosphate binding | molecular_function |
| GO:0070936 | Rnf216 | missense_variant | involved_in | protein K48-linked ubiquitination | biological_process |
| GO:0097746 | Cbs | missense_variant | acts_upstream_of_or_within | blood vessel diameter maintenance | biological_process |
| GO:0098685 | Rnf216 | missense_variant | located_in | Schaffer collateral - CA1 synapse | cellular_component |
| GO:0098685 | Rnf216 | missense_variant | is_active_in | Schaffer collateral - CA1 synapse | cellular_component |
| GO:0098690 | Glra4 | missense_variant | located_in | glycinergic synapse | cellular_component |
| GO:0098843 | Rnf216 | missense_variant | located_in | postsynaptic endocytic zone | cellular_component |
| GO:0098978 | Homer2 | missense_variant | is_active_in | glutamatergic synapse | cellular_component |
| GO:0098978 | Homer2 | missense_variant | located_in | glutamatergic synapse | cellular_component |
| GO:0098978 | Rnf216 | missense_variant | located_in | glutamatergic synapse | cellular_component |
| GO:0099060 | Glra4 | missense_variant | located_in | integral component of postsynaptic specialization membrane | cellular_component |
| GO:1902476 | Glra4 | missense_variant | involved_in | chloride transmembrane transport | biological_process |
| GO:1902476 | Glra4 | missense_variant | involved_in | chloride transmembrane transport | biological_process |
| GO:1990837 | Gcm1 | missense_variant | enables | sequence-specific double-stranded DNA binding | molecular_function |
