## Supplemental Table 2 for "Divergent risky decision-making and impulsivity behaviors in Lewis rat substrains with low genetic difference"

Table S2. Association of genes with predicted high to moderate impact variants with human psychiatric diseases

| **Gene or Intergenic region** | **Mapped Trait** | **PMID** |
| --- | --- | --- |
| BEND4 | helping behavior measurement | 29324852 |
| BEND4 | unipolar depression, autism spectrum disorder | 30804558 |
| CEACAM3 | smoking behavior, 3-hydroxy-1-methylpropylmercapturic acid measurement | 26053186 |
| CGAS | smoking behavior, 3-hydroxy-1-methylpropylmercapturic acid measurement | 26053186 |
| CGAS | smoking behavior, 3-hydroxypropylmercapturic acid measurement | 26053186 |
| CTU2 | autism spectrum disorder symptom | 25534755 |
| GLIS3 | schizophrenia | 26198764 |
| GRIN2B | nicotine dependence symptom count, depressive symptom measurement | 30287806 |
| GRIN2B | schizophrenia | 26198764 |
| GRIN2B | risk tolerance | 30643258 |
| GRIN2B | unipolar depression | 23377640 |
| GRIN2B | unipolar depression, response to selective serotonin reuptake inhibitor | 30034349 |
| Hip1r | schizophrenia | 30285260 |
| Hip1r | schizophrenia | 28991256 |
| Hip1r | schizophrenia | 31740837 |
| IP6K3 | schizophrenia, intelligence, self reported educational attainment | 31374203 |
| LIN28B | schizophrenia | 26198764 |
| LIN28B | schizophrenia | 28991256 |
| LIN28B | schizophrenia | 30285260 |
| LIN28B | schizophrenia | 31740837 |
| LIN28B | smoking behavior, BMI-adjusted waist circumference | 28443625 |
| LIN28B | unipolar depression | 27479909 |
| MAP2K5 | bipolar disorder, body mass index | 31754094 |
| MAP2K5 | smoking behavior | 30643258 |
| MAP2K5 | smoking behavior, body mass index | 28443625 |
| Map2k5 | bipolar disorder | 31754094 |
| MAP3K20 | smoking behavior | 30643258 |
| MSI2 | schizophrenia | 26198764 |
| MSI2 | schizophrenia | 28991256 |
| MSI2 | schizophrenia | 30285260 |
| MSI2 | schizophrenia | 31740837 |
| NIN | schizophrenia | 33169155 |
| NRG3 | schizophrenia, antipsychotic drug use measurement, schizoaffective disorder | 26821981 |
| POC1B | schizophrenia | 26198764 |
| RGS6 | anorexia nervosa, obsessive-compulsive disorder, attention deficit hyperactivity disorder, Tourette syndrome, unipolar depression, schizophrenia, autism spectrum disorder, bipolar disorder | 31835028 |
| RGS6 | bipolar disorder | 32606422 |
| RGS6 | schizophrenia | 25056061 |
| RGS6 | schizophrenia | 26198764 |
| RGS6 | schizophrenia | 28991256 |
| RGS6 | schizophrenia | 29483656 |
| RGS6 | schizophrenia | 30285260 |
| RGS6 | schizophrenia | 31268507 |
| RGS6 | schizophrenia | 31740837 |
| RGS6 | schizophrenia | 32606422 |
| RGS6 | schizophrenia, antipsychotic drug use measurement, schizoaffective disorder | 26821981 |
| RGS6 | schizophrenia, autism spectrum disorder | 28540026 |
| RGS6 | schizophrenia, intelligence, self reported educational attainment | 31374203 |
| SBK1 | smoking behavior, body mass index | 28443625 |
| SLC12A5 | unipolar depression | 30718901 |
