## Supplemental Table 3 for "Divergent risky decision-making and impulsivity behaviors in Lewis rat substrains with low genetic difference"

**Table S3.** Whole genome sequencing data of inbred rats from NIH SRA

| **Strain** | **SRA Run IDs** |
| --- | --- |
| ACI/EurMcwi | ERR224446 |
| ACI/N | SRR7755593 |
| BBDP/Wor | ERR224447 |
| BN/N | SRR7755594 |
| BN/NHsdMcwi | ERR5309015 ERR5309016 ERR5309017 ERR5309018 ERR5309019 ERR5309020 ERR5309021 ERR5309022 |
| BN/NHsdMcwi | SRR14880436 |
| BN/NHsdMcwi | SRR14880437 |
| BN/NHsdMcwi | SRR14880438 |
| BN/NHsdMcwi | SRR14880439 |
| BUF/N | SRR7755597 |
| DA | SRR351198 SRR351199 SRR351207 SRR351208 SRR351209 SRR351210 SRR351211 SRR351212 |
| F344 | SRR351197 SRR351200 SRR351201 SRR351202 SRR351203 SRR351204 SRR351205 SRR351206 |
| F344/N | SRR7755598 |
| F344/NCrl | ERR224448 |
| FHH/EurMcwi | ERR224449 |
| FHL/EurMcwi | ERR224450 |
| GK/Ox | ERR224451 |
| HSRA/Mg | SRR10233451 |
| HSRA/Mg | SRR10233452 |
| LE/Stm | ERR224452 |
| LEW/Crl | ERR224453 |
| LEW/NCrlBr | ERR224454 |
| LH/MavRrrc | ERR224455 |
| LL/MavRrrc | ERR224457 |
| LN/MavRrrc | ERR224456 |
| M520/N | SRR7755599 |
| MHS/Gib | ERR224458 |
| MNS/Gib | ERR224459 |
| MR/N | SRR7755600 |
| SBH/Ygl | ERR224460 |
| SBN/Ygl | ERR224461 |
| SHR/NHsd | ERR224462 |
| SHR/SPGla | ERR224463 |
| SR/Jr | ERR224464 |
| SS/Jr | ERR224465 |
| SS/JrHsdMcwi | ERR224466 |
| WAG/Rij | ERR224467 |
| WKY/Gla | ERR224469 |
| WKY/N | SRR7755595 |
| WKY/NCrl | ERR224468 |
| WKY/NHsd | ERR224470 |
| WN/N | SRR7755596 |
