## Supplemental Figure Captions for "Divergent risky decision-making and impulsivity behaviors in Lewis rat substrains with low genetic difference"

**Figure S1. Comparing sequencing data from Lewis substrains and other inbred strains on chromosome 8**

a. number of variants, b. read depth, c. genotype quality, and d. number of missing calls. The 78.0-79.5 Mb region, where most of the variants unique to each Lewis substrain on chr8 were located, was marked by vertical lines. The “other” samples were the mean of 36 samples downloaded from NCBI SRA (see Table S3).

**Figure S2. Matrix view of chromosome 8 between 79 - 79.3 Mb**

a. BN/NHsdMcwi b. Lew/NCrl, and c. Lew/NHsd. The matrix view was constructed from data on the overlap of sequencing barcodes used in the linked-read libraries. These overlaps could indicate complex structural variants, such as those identified by the blue boxes.

**Figure S3. Phylogenetic analysis of Chromosome 8 using data from 41 rat samples**

a. Entire chromosome 8. b. Genomic segment between 78.6-79.3 Mb.

**Figure S4. Genetic variants shared between Lewis substrains and related strains**
