## Supplementary figures and images for "Divergent risky decision-making and impulsivity behaviors in Lewis rat substrains with low genetic difference"

### Supplemental Figure 1

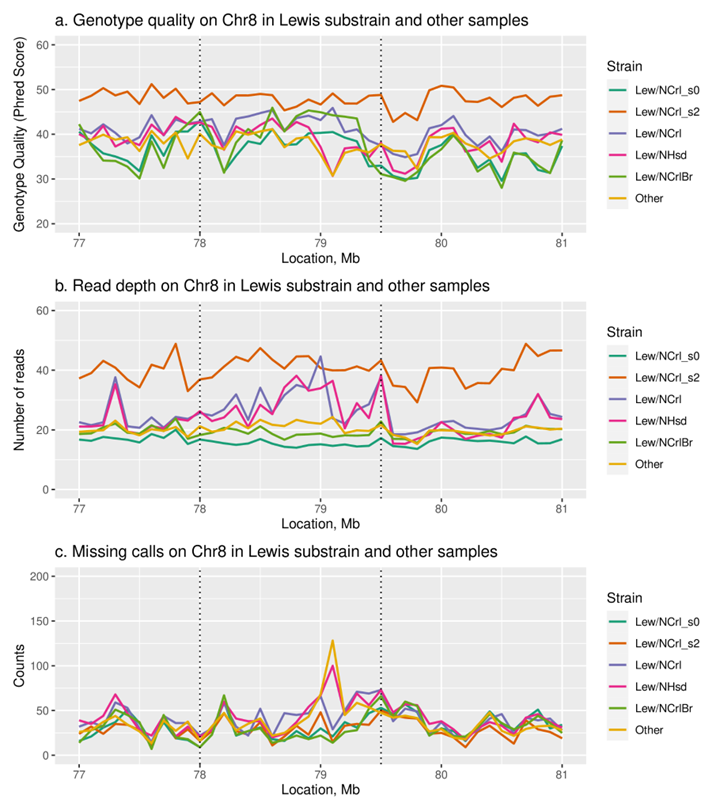

### Supplemental Figure 2

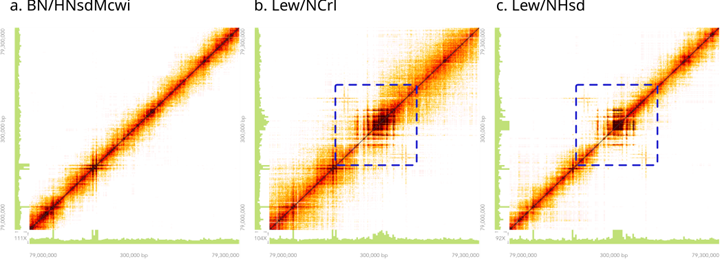

### Supplemental Figure 3

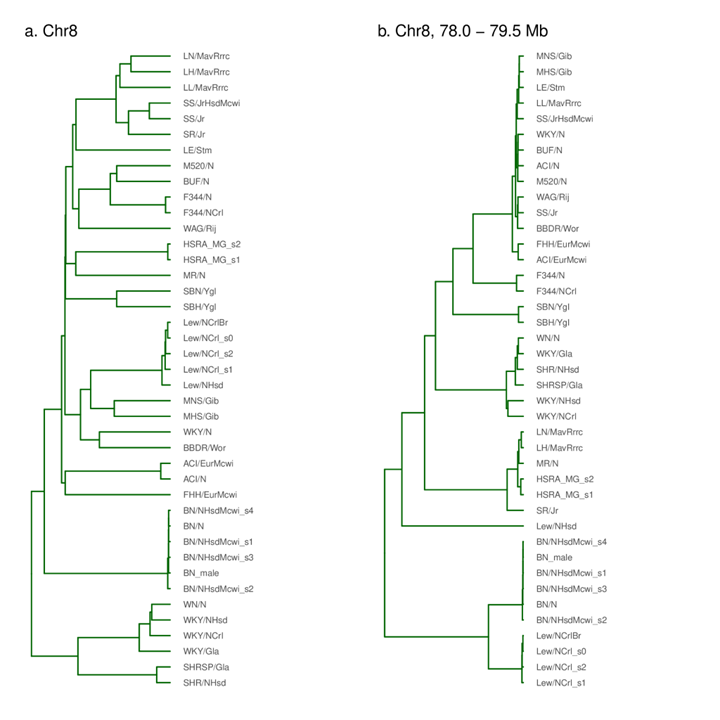

### Supplemental Figure 4

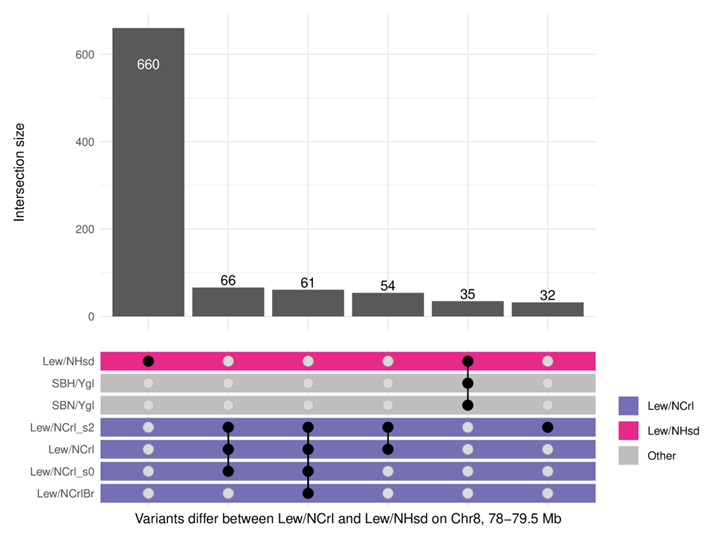
